# Sprint interval exercise disrupts mitochondrial ultrastructure driving a unique mitochondrial stress response and remodelling in humans

**DOI:** 10.1101/2024.12.19.629456

**Authors:** J. Botella, E. Perri, N. J. Caruana, S. López-Calcerrada, M. Brischigliaro, N. A. Jamnick, V. Oorschot, N. J. Saner, J. Diaz-Lara, D. F. Taylor, A. Garnham, E. Fernández-Vizarra, C. Ugalde, G. Ramm, D. A. Stroud, M. Lazarou, D. J. Bishop

**Affiliations:** Institute for Health and Sport (IHES), Victoria University, Melbourne, Australia; Metabolic Research Unit, School of Medicine and Institute for Mental and Physical Health and Clinical Translation (IMPACT), Deakin University, Waurn Ponds, Australia; Department of Biomedical Sciences for Health, University of Milan, Italy; Department of Biochemistry and Pharmacology, Bio21 Molecular Science and Biotechnology Institute, The University of Melbourne, Melbourne, Australia; Hospital 12 de Octubre Research Institute (i+12), 28041 Madrid, Spain; Veneto Institute of Molecular Medicine, 35131 Padova, Italy; Department of Biomedical Sciences, University of Padova, 35131 Padova, Italy; Department of Neurology, University of Miami Miller School of Medicine. Miami, FL 33136, USA; Monash Ramaciotti Centre for Cryo Electron Microscopy, Monash University, Melbourne, Australia; Electron Microscopy Core Facility, European Molecular Biology Laboratory, Heidelberg, Germany; Performance and Sport Rehabilitation Laboratory, Faculty of Sports Sciences, University of Castilla-La Mancha, 45071, Toledo, Spain; Department of Biochemistry and Molecular and Cellular Biology, Faculty of Health and Sport Sciences, University of Zaragoza, 22002 Huesca, Spain; Margarita Salas Center for Biological Research (CIB-CSIC), Madrid, Spain; Centro de Investigación Biomédica en Red de Enfermedades Raras (CIBERER), U723, Madrid, Spain; Murdoch Children’s Research Institute, Royal Children’s Hospital, Parkville, VIC, Australia; Victorian Clinical Genetics Services, Royal Children’s Hospital, Parkville, VIC, Australia; Department of Biochemistry and Molecular Biology, Biomedicine Discovery Institute, Monash University, Melbourne, Australia; Walter and Eliza Hall Institute of Medical Research, Melbourne, Australia; Department of Medical Biology, University of Melbourne, Melbourne, Australia

**Keywords:** skeletal muscle, mitochondria, exercise, UPR^mt^, mitophagy

## Abstract

Exercise remains the most effective lifestyle intervention to remodel the mitochondrial network and to prevent most non-communicable diseases. Despite this, the molecular mechanisms by which different exercise prescriptions dictate mitochondrial remodelling are poorly understood in humans. Here, we show that, compared to moderate-intensity continuous exercise (MICE), sprint-interval exercise (SIE) – a known time-efficient high-intensity exercise – leads to mitochondrial stress and activates the mitochondrial unfolded protein response (UPR^mt^). The SIE-specific signature is characterized by a morphological and ultrastructural mitochondrial disturbance, concurrent with the activation of the integrated stress response (ISR) and mitochondrial quality control (MQC) pathways. When the respective exercises are repeated over time (8 weeks), our results demonstrate that moderate-intensity continuous training (MICT) and sprint-interval training (SIT) lead to a divergent mitochondrial remodelling. MICT elicits a mitochondrial adaptation characterized by an increase in markers of mitochondrial content, complex I activity, and enrichment of proteins involved in tricarboxylic acid (TCA) cycle and oxidative phosphorylation (OXPHOS) system. On the other hand, SIT leads to proteomic enrichment of pathways involved in mitochondrial 1-Carbon metabolism and protein quality control, concurrently with improvements in mitochondrial respiratory function. Lastly, we have identified COX7A2L as a divergently regulated protein across groups, significantly accumulating in III_2_+IV_1_ respiratory supercomplexes only following SIT. In conclusion, our study provides mechanistic insights on how SIE and MICE divergently impact the post-exercise mitochondrial signalling, and subsequent long-term mitochondrial remodelling following training. These findings provide a strong basis for targeted exercise prescription to modulate specific mitochondrial adaptations in human skeletal muscle.

## INTRODUCTION

Mitochondria are critical organelles involved in many biological functions, including metabolism, inflammation, and cell death (Spinelli and Haigis, 2018). Owing to this, suboptimal mitochondrial characteristics (e.g., decreased respiratory function) have been implicated in various medical conditions (Grevendonk et al., 2021; Migliavacca et al., 2019). These same mitochondrial characteristics are often enhanced in the skeletal muscle of athletes (Botella et al., 2023a; Hoppeler et al., 1973; Jacobs and Lundby, 2013; Nielsen et al., 2017). Given the large energetic demands of contracting skeletal muscle (Sahlin et al., 1998), it is not surprising that exercise provides a powerful stimulus that can trigger enduring changes to mitochondrial content, enzyme activity, and respiratory chain function.

Exercise is a well-known intervention that stimulates the synthesis of new mitochondrial components – a process termed mitochondrial biogenesis (Botella et al., 2022; Holloszy, 1967). While many of the underlying mechanisms are poorly understood (Neufer et al., 2015; Zierath and Wallberg-Henriksson, 2015), exercise also remains one of the most effective lifestyle interventions for the prevention of non-communicable diseases (Booth et al., 2012; Lee et al., 2012). The benefits of exercise are thought to stem from the contraction-induced homeostatic disturbances that initiate an orchestrated change in the molecular landscape (i.e., post-translational modifications, gene expression, etc) aiming to re-establish cellular homeostasis (Blazev et al., 2022; Perry et al., 2010). This response is largely dependent on the characteristics of the exercise (Hood et al., 2011; Robinson et al., 2017). Elucidating the cellular programs affected by distinct exercise prescriptions is important to facilitate the establishment of more evidence-based exercise guidelines and targeted exercise interventions to improve health (Bishop et al., 2024).

High-intensity interval exercise (HIIE) has gained popularity in the last decades due to its effectiveness in stimulating skeletal muscle and cardiorespiratory adaptations traditionally associated with moderate-intensity continuous exercise (MICE) (Burgomaster et al., 2008; Cochran et al., 2014; Gillen et al., 2016). The most time-efficient form of HIIE is termed sprint-interval exercise (SIE) and requires as little as four to six 30-second all-out sprints (Gibala and Hawley, 2017), making it an attractive exercise prescription given the time constraints of modern society (Bartlett et al., 2011; Godin et al., 1994). While it remains a topic of debate, multiple mechanisms have been proposed to underlie the efficiency of SIE. Some have shown that SIE is associated with a stress response that leads to cellular calcium leakage and the subsequent activation of pathways involved in mitochondrial biogenesis (Place et al., 2015; Zanou et al., 2021). Others have reported an increase in the abundance of nucleus-localized transcription factors implicated in mitochondrial biogenesis (e.g., peroxisome proliferator activated receptor δ coactivator 1α; PGC-1α) following SIE but not MICE (Granata et al., 2017). Many studies have suggested this greater activation of mitochondrial biogenesis as one of the underlying mechanisms contributing to the efficiency of SIE. However, the paradoxical observation that SIE, when repeated over time (i.e., in exercise training), does not consistently translate into larger increases in mitochondrial content adaptations (Bishop et al., 2019b; Granata et al., 2018; Granata et al., 2016a) suggests that other, yet-to-be discovered, mechanisms may contribute to the efficacy of SIE.

Exercise has been hypothesised to promote a mitochondrial hormetic (termed mitohormesis) response to re-establish mitochondrial homeostasis, leading to a better tolerance to subsequent stressors (Merry and Ristow, 2016). Multiple pathways, such as those controlling mitochondrial dynamics (fusion, fission), mitochondrial quality control (MQC; e.g., mitophagy), and the mitochondrial unfolded protein response (UPR^mt^), are known to mediate the maintenance of a healthy mitochondrial pool (Melber and Haynes, 2018; Shpilka and Haynes, 2018). In mice, exercise has been shown to influence most of these pathways (Cordeiro et al., 2020; Cordeiro et al., 2021; Laker et al., 2017; Lavorato et al., 2018; Picard et al., 2013) but there is limited understanding of the influence of exercise on these pathways in humans (Houzelle et al., 2021; Huertas et al., 2019) and how these pathways are modified by different exercise prescriptions. Given the larger homeostatic disturbances associated with high-intensity exercise, a plausible hypothesis is that mitochondrial stress is an important mechanism contributing to the mitochondrial adaptations seen in SIE.

Here, we report that SIE, but not MICE, disturbs the mitochondrial ultrastructure leading to mitochondrial stress and the transcriptional activation of the UPR^mt^ in human skeletal muscle. By combining an array of complementary techniques (e.g., RNA sequencing, electron microscopy, proteomics), we provide strong evidence of a SIE-specific mitochondrial stress and a concurrent upregulation of the integrated stress response (ISR) and MQC pathways - largely independent of markers of mitochondrial biogenesis and dynamics. When repeated over time (8 weeks), only moderate-intensity continuous training (MICT) increased markers of mitochondrial content, while only sprint-interval training (SIT) increased mitochondrial respiratory chain function. Guided by our proteomics approach, we observe that SIT divergently remodels the mitochondrial proteome and upregulates the expression of COX7A2L protein and its incorporation into III_2_IV_1_ respiratory supercomplexes, providing a novel mechanism contributing to the effectiveness of SIT. These findings challenge our current knowledge of exercise-specific mitochondrial adaptations, while also supporting the importance of personalized training interventions to achieve specific mitochondrial remodelling.

## RESULTS

### There are marked physiological differences between moderate-intensity continuous exercise (MICE) and sprint-interval exercise (SIE)

Improvements in various mitochondrial characteristics are a hallmark of exercise training (Bishop et al., 2019a; Hood et al., 2019). While distinct mitochondrial adaptations have been linked to the intensity and volume of the exercise prescribed (Granata *et al*., 2018), it remains a topic of debate (Bishop *et al*., 2019b; MacInnis et al., 2019). In the current study, we recruited twenty-eight healthy males to uncover the divergent effects of exercise intensity and volume (Table 1 and Figure 1A). We chose SIE as it is the most intense type of HIIE and is widely recognized as an effective and time-efficient exercise (Gibala and Hawley, 2017). To study the effects of exercise volume we prescribed MICE, which is generally viewed as the ‘traditional’ type of endurance or cardiorespiratory exercise (Garber et al., 2011). Participants were assigned in a random counter-balanced order to one of the two exercise groups based on their VLO_2max_ and WL_max_ (Table 1). A *posteriori* analysis showed that both groups were matched for age, as well as for anthropometrical parameters, and markers of mitochondrial content and respiratory function (all *p* > 0.05; Table 1 and Data S1).

**Figure 1.**
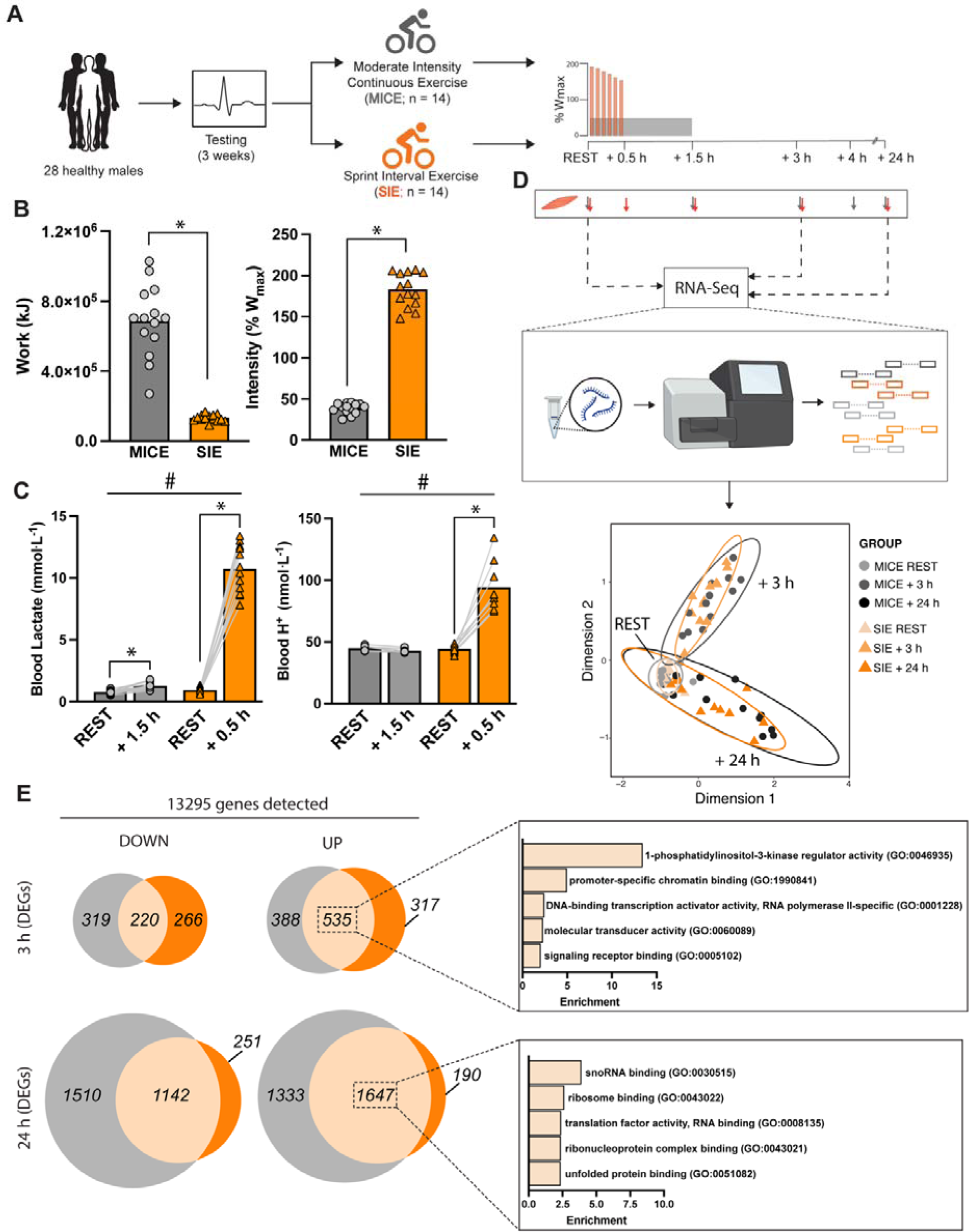
Schematic of the experimental exercise session design and exercise characteristics. A) Workflow of the study including participant recruitment, testing, and allocation to either moderate-intensity continuous exercise (MICE) or sprint-interval exercise (SIE). B) Exercise volume and intensity achieved during the experimental exercise session for both MICE (n = 14) and SIE (n = 14). C) Exercise-induced changes in blood lactate and hydrogen (H+) concentrations for both MICE (n = 14 for lactate, n = 7 for H+) and SIE (n = 14 for lactate, n = 9 for H+)). D) Selected muscle biopsy timepoints (grey arrows – MICE biopsy timepoint; orange arrows – SIE biopsy timepoints; n = 12 per group) for RNA-sequencing and their distribution via principal component analysis (PCA) showing a largely shared clustering. E) Venn diagrams showing the number of common and unique differentially expressed genes (DEGs) at each timepoint. Overrepresentation analysis (ORA) of the molecular functions commonly regulated by both types of exercise at 3 hours (early recovery) and at 24 hours (late recovery). * denotes p < 0.05 between groups or timepoints; # denotes time x group interaction of p < 0.05. Data are expressed as mean. For B) differences were assessed via one-way ANOVA. For C) differences were assessed via two-way ANOVA with Sidak’s multiple comparison test.

**Table 1.** Baseline characteristics of the participants in the two exercise training groups.

|  | **MICE/MICT**<br>**(n=14)** | **SIE/SIT**<br>**(n=14)** | **ANOVA *p* value** |
| --- | --- | --- | --- |
| **Biological** |  |  |  |
| Age (years) | 26.1 ± 5.7 | 26.9 ± 4.9 | 0.685 |
| Body mass (kg) | 77.0 ± 10.7 | 76.5 ± 9.7 | 0.909 |
| BMI (kg·m<sup>-2</sup>) | 24.1 ± 2.6 | 23.7 ± 2.8 | 0.748 |
| **Physiological** |  |  |  |
| V̇O<sub>2max</sub> (mL·min<sup>-1</sup>·kg<sup>-1</sup>) | 52.2 ± 6.4 | 51.9 ± 6.5 | 0.789 |
| Ẇ<sub>max</sub> (W·kg<sup>-1</sup>) | 4.2 ± 0.6 | 4.2 ± 0.8 | 0.998 |
| 20km-TT (Ẇ·kg<sup>-1</sup>) | 2.7 ± 0.5 | 2.7 ± 0.5 | 0.955 |
| 4km-TT (Ẇ·kg<sup>-1</sup>) | 3.2 ± 0.5<br>(n = 13) | 3.1 ± 0.5 | 0.741 |
| **Molecular** |  |  |  |
| Mitochondrial volume density (%) | 5.2 ± 0.7<br>(n = 10) | 5.4 ± 1.6<br>(n = 12) | 0.728 |
| Citrate Synthase Activity (mol·h<sup>-1</sup>·kg protein<sup>-1</sup>) | 2.4 ± 0.5<br>(n = 12) | 2.6 ± 0.7<br>(n = 12) | 0.459 |
| Mitochondrial respiratory function (ETF+CI+IIp; pmol O<sub>2</sub>·s<sup>-1</sup>·mg protein<sup>-1</sup>) | 83.2 ± 13.7<br>(n = 9) | 83.8 ± 23.4<br>(n = 10) | 0.952 |

MICE = Moderate-intensity continuous exercise; MICT = Moderate-intensity continuous training; SIE = sprint interval exercise; SIT = sprint interval training; BMI = body mass index; V̇O<sub>2max</sub> = maximal rate of oxygen consumption; Ẇ<sub>max</sub> = peak aerobic power; 20km-TT = 20-kilometre time trial; 4km- TT = 4-kilometre time-trial. ETF+CI+IIp = Maximal respiration through electron-transferring flavoprotein, complex I, and complex II (ADP-stimulated in the presence of pyruvate and succinate). Data are expressed as mean ± SD. Differences between groups were assessed via one-way ANOVA.

Following exercise familiarization, testing, and standardization of macronutrient intake, participants completed their assigned experimental trial where *vastus lateralis* skeletal muscle biopsies were collected (Figure 1A and S1A). While a work-matched approach is often used to study between-exercise differences, this was not possible as a greater exercise volume inherently implies a larger amount of work completed. Therefore, to emphasise the distinguishing characteristics of each exercise, we designed the exercises to have similar between-group differences for intensity and volume (Figure 1B). Participants conducting MICE completed a ∼ 5.2-fold higher exercise volume, while participants performing SIE exercised at a ∼ 4.7-fold higher intensity. Both exercises elicited an increase in blood lactate concentration (Figure 1C), with a larger increase following SIE than MICE (∼10-fold vs ∼2-fold; *p* < 0.05). This greater increase in lactate following SIE is in line with previous evidence indicating that exercise intensity is a driver of increased glycolytic flux and lactate production during exercise (Jamnick et al., 2020; van Loon et al., 2001). Concurrently, only SIE, but not MICE, led to an increase in blood H^+^ concentration (*p* < 0.05; Figure 1C) and elicited a greater rating of perceived exertion (*p* < 0.05; Figure S1B). The larger metabolic disturbance elicited by SIE compared with MICE is consistent with previous findings (Granata *et al*., 2017; Parker et al., 2017).

### Transcriptome divergence is observed between MICE and SIE in the early recovery from exercise

The physiological adaptations to exercise are thought to partially stem from an orchestrated transcriptional response to the exercise stimulus (Perry *et al*., 2010; Pillon et al., 2020). Given the very different metabolic and physiological responses between MICE and SIE, we hypothesized there would also be divergent transcriptional responses to these two exercises. We performed RNA-sequencing analyses on the resting muscles samples (rest) and at the early (+ 3 h) or late (+ 24 h) recovery points from the start of exercise (Figure 1D). We detected 13295 gene transcripts across all samples, with 7175 transcripts differentially expressed in response to at least one exercise stimuli and at least at one timepoint (representing ∼ 54% of the detected transcriptome). This far exceeds the number of transcripts that were identified as being altered by exercise identified in two previous meta-analyses (Amar et al., 2021; Pillon *et al*., 2020), possibly due to the greater standardization possible with our parallel-group study design and the inclusion of the 24 hour biopsy time. Our data highlights that many of the changes in the exercise-induced transcriptome have remained ‘hidden’ because of most studies only taking biopsies in the first few hours post exercise.

In the early recovery from exercise there were 1462 and 1338 transcripts that were differentially expressed following MICE and SIE, respectively. In the late recovery, there was a larger number of transcripts altered with 5632 and 3230 differentially expressed transcripts following MICE and SIE, respectively (Figure 1E). While a previous meta-analysis identified 159 transcripts that followed specific post-exercise trajectories (irrespective of exercise mode; (Amar *et al*., 2021)), our results have captured a much larger temporal effect. These findings provide an important resource that helps characterize the time-dependent complexity of the transcriptional responses to exercise (Amar *et al*., 2021; Kuang et al., 2022).

Across exercise groups, there were 755 and 2789 shared differentially expressed transcripts in the early and late recovery phase, respectively (Figure 1E and Data S1). The transcripts commonly regulated by both groups, which point to the shared transcriptional response to endurance exercise, demonstrated an overrepresentation of a broad range of cellular pathways differently altered early (+ 3h) and late in the recovery (+ 24h) (Figure 1E and Data S1). While our experimental design allowed us to establish a common early and late transcriptomic signature, future studies should explore a more extensive exercise-induced transcriptional time course with the contribution of diverse cell types to these transcriptional changes (Lovrić et al., 2022).

### The mitochondrial unfolded protein response (UPR^mt^) is a distinct transcriptional signature following sprint interval exercise (SIE)

Having established the existence of common transcriptional changes following different types of endurance exercise, we next aimed to uncover the transcriptional differences between MICE and SIE. These results could point toward the molecular signalling pathways that may underlie some of the beneficial adaptations previously reported in response to SIE (Gibala and Hawley, 2017). Despite the larger number of altered transcripts observed in the late recovery (i.e., 24 h; Figure 1E and Data S1), there were only 16 significant differentially-regulated transcripts between MICE and SIE at this timepoint (Adj. p < 0.05; Data S2). There was a greater between-group divergence early in the recovery with 207 transcripts differentially regulated between MICE and SIE 3 h post-exercise (Data S2). We therefore focused our attention on the early timepoint (i.e., 3 h) and performed hierarchical clustering of these 207 transcripts to further interrogate the underlying biological processes associated with these transcripts (Figure 2A). Of these differences, 116 were significantly upregulated to a larger extent following SIE (and 91 were upregulated to a larger extent following MICE). By performing overrepresentation analyses (ORA), significant enrichments were only observed for the list of genes upregulated to a larger extent following SIE. These transcripts included numerous cytosolic and mitochondrial heat-shock proteins (HSPs) and showed a systematic enrichment for terms related to protein folding and the unfolded protein response (UPR) (Figure 2A and 2B).

**Figure 2.**
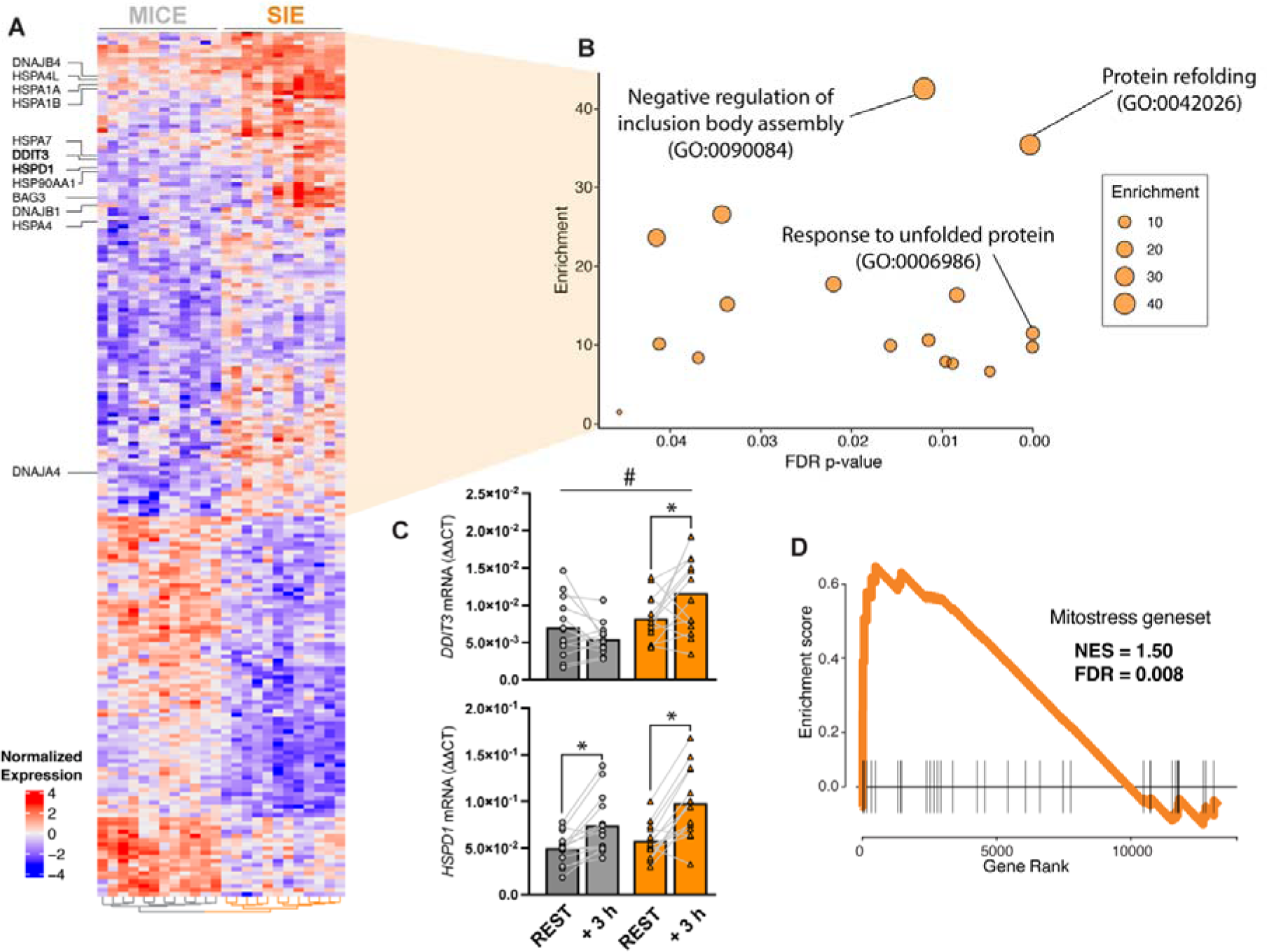
Transcriptomic divergence in the early recovery of MICE and SIE. A) Hierarchical clustering of the divergently regulated transcripts at 3 hours following both moderate-intensity continuous exercise (MICE) and sprint-interval exercise (SIE). Selected transcripts known to be involved in the unfolded protein response (UPR) are listed on the left of the heat map. B) Bubble plot of the pathways positively enriched only following SIE using overrepresentation (ORA) analysis. C) mRNA levels of DDIT3 and HSPD1 - key transcripts involved in the mitochondrial UPR (UPR^mt^) and identified via RNAseq as being upregulated to a greater extent following SIE (n = 14) vs MICE (n = 13). D) Gene set enrichment analysis (GSEA) of a previously identified mitochondrial stress geneset (termed ‘mitostress’ from Quiros et al., 2017), which shows an enrichment only following SIE exercise. NES = Normalized Enrichment Score. FDR = False Discovery Rate. GO = Gene Ontology. * denotes p < 0.05 between timepoints; # denotes time x group interaction is p < 0.05. Data are expressed as mean. For C) differences between groups were assessed via 2-way ANOVA with Sidak’s multiple comparison test and pre-planned within-group one-way ANOVA.

Two of the differentially expressed genes between SIE and MICE were DNA Damage Inducible Transcript 3 (*DDIT3*; also known as CHOP) and Heat Shock Protein Family D Member 1 (*HSPD1*; also known as HSP60) - both of which were the first two identified components of the mammalian UPR^mt^ (Zhao et al., 2002). These findings were subsequently validated via qPCR (Figure 2C and Figure S2A), and it was further observed that the mitochondrial protease *HSPE1*, also involved in the UPR^mt^, was increased to a larger extent 3 h following SIE (Figure S2A). These results pointed towards a coordinated upregulation of mitochondrial and cytosolic HSPs following SIE, which is consistent with the suggestion that the UPR^mt^ transcriptional response requires the coordinative activation of both cytosolic and mitochondrial HSPs (Sutandy et al., 2023; Wrobel et al., 2015).

To confirm the mitochondrial origin of the stress following SIE, we probed our datasets against the ‘mitostress’ geneset - composed of genes previously shown to be commonly upregulated following different *in vitro* mitochondrial stressors (Quiros et al., 2017). In accordance with our findings above, the ‘mitostress’ geneset was significantly enriched only following SIE (normalized enrichment score (NES) = 1.50; False discovery rate (FDR) < 0.01; Figure 2D). The upregulation of the ‘mitostress’ geneset only occurred in the early recovery from SIE (Figure S2B), suggesting that the mitochondrial stress and UPR^mt^ response occurred early following SIE, and had dissipated 24 hours into the recovery – indicative of SIE being a mitohormetic stressor (i.e., reversible and adaptive) (Sprenger and Langer, 2019; Yun and Finkel, 2014).

### Sprint exercise elicits acute mitochondrial ultrastructural and morphological disturbances

To better understand the events preceding the transcriptional upregulation of the UPR^mt^, we next used transmission electron microscopy (TEM) to quantify morphological and ultrastructural changes to mitochondria before and after both MICE and SIE (Figure 3A). Across all morphological variables, there were no significant changes following MICE (Figure 3B; Data S3), which is consistent with the absence of the mitochondrial stress signature following this type of exercise in our RNA-seq data (Figure S2B). In contrast, following SIE, there was a significant increase in the roundness of mitochondria, with a concurrent decrease in the aspect ratio (indicating a decrease in mitochondrial complexity; Figure 3B). We also observed an increase in mitochondrial size following SIE (measured as area; Figure 3B), which suggested swollen mitochondria (Wang et al., 2019) and the presence of mitochondrial stress (Wai and Langer, 2016), in accordance with our RNA-seq data.

**Figure 3.**
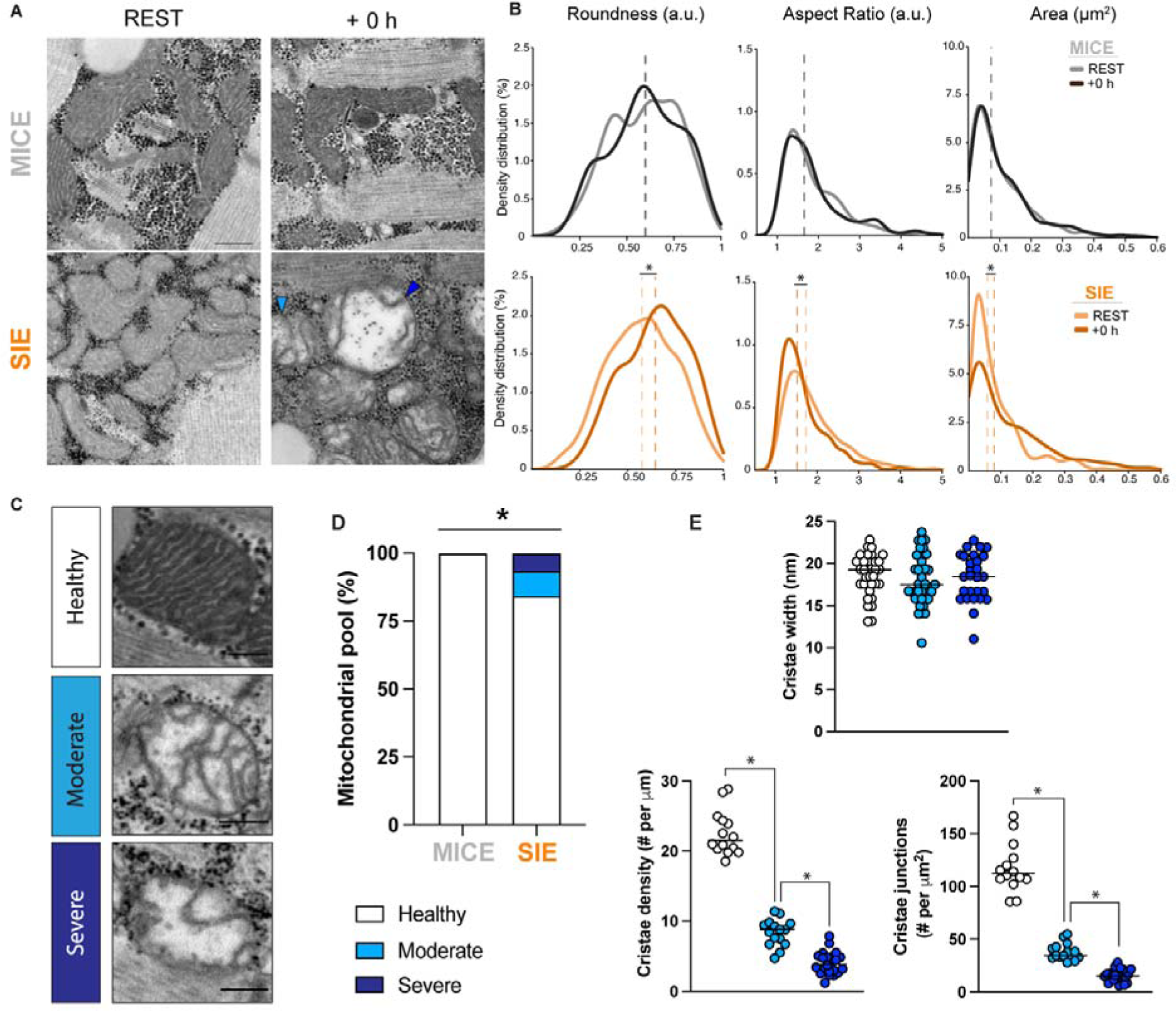
Morphological and ultrastructural disturbances to mitochondria following SIE. A) Representative transmission electron microscopy (TEM) images of skeletal muscle before and after moderate-intensity continuous exercise (MICE; n = 4) and sprint-interval exercise (SIE; n = 5). The light blue arrow points to a moderately damaged mitochondrion, while the dark blue arrow points to a severely damaged mitochondrion. There were no damaged mitochondria identified in the samples obtained at +0 h following MICE. Scale bar = 0.5 µm. B) Density distribution of the area, roundness, and aspect ratio of mitochondrial profiles before and after MICE (REST, n = 428; + 0 h, n = 446) and SIE (REST, n = 603; + 0 h, n = 551). C) Representative mitochondria from each category of healthy, moderate, or severe damage. Scale bar = 0.2 µm. D) Distribution of mitochondrial profiles identified as healthy, and moderate or severely damaged following both MICE and SIE. E) Mitochondrial cristae width, density, and number of junctions across representative mitochondria from each subgroup. * denotes group difference at p < 0.05. Data are expressed as mean. For D) differences between groups were assessed via one-way ANOVA. For E) one-way ANOVA with Sidak’s multiple comparisons test.

Given the clear transcriptional and morphological mitochondrial stress signature observed following SIE, we next performed a qualitative analysis to detect any mitochondrial ultrastructural disturbances. Strikingly, we observed that following SIE, on average, 16% of the analyzed mitochondrial pool had signs of ultrastructural disturbance (Figure 3C and 3D); this phenomenon was not observed at rest in either group or following MICE. These mitochondrial disturbances, further categorized into moderate or severe, were associated with decreased number of cristae density and number of cristae junctions, but not changes in cristae width (Figure 3E and Data S3). Similar mitochondrial ultrastructural disturbances have previously been reported following endurance exercise in non-human skeletal muscle (running to exhaustion in rats (Gollnick and King, 1969) and long-duration running or sprinting in horses (Nimmo and Snow, 1982)), but were not previously observed in humans following moderate-intensity endurance exercise (Gollnick and King, 1971) - in accordance with our own results following MICE. The present results are, to the best of our knowledge, the first to report disturbances in mitochondrial ultrastructure following endurance exercise in humans. Disruptions to mitochondrial ultrastructure would presumably alter processes dependent on the membrane potential, such as protein import and ATP synthesis (Glancy et al., 2017; Wolf et al., 2019). Collectively, these results corroborate our transcriptomic results and point towards a new paradigm whereby SIE acutely provokes mitochondrial ultrastructural disturbances in human skeletal muscle and initiates a mitochondrial-specific stress response.

### The integrated stress response (ISR) and mitochondrial quality control (MQC) pathways are upregulated following SIE

Given our transcriptomic (RNA-seq) and electron microscopy (TEM) findings, we hypothesized that pathways associated with mitochondrial stress would be differentially activated by the two types of exercise tested. In humans, mitochondrial stress signals through the integrated stress response (ISR) to restore cellular homeostasis (Fessler et al., 2020; Guo et al., 2020; Pakos-Zebrucka et al., 2016; Quiros *et al*., 2017). Therefore, we quantified the phosphorylation of the ISR factor Eukaryotic translation initiation factor 2A (eIF2α) and the mRNA expression of Protein Phosphatase 1 Regulatory Subunit 15A (*PPP1R15A*; also known as GADD34), which are both increased following ISR activation (Pakos-Zebrucka *et al*., 2016). In support of our hypothesis, we observed a larger increase in eIF2α phosphorylation and *PPP1R15A* mRNA expression following SIE than after MICE (Figure 4A). Similarly, we observed a larger increase following SIE of the activating transcription factor 3 (*ATF3*) mRNA and fibroblast growth factor 21 (*FGF21*) mRNA (Figure S3A), both known to be upregulated following chronic mitochondrial stress in skeletal muscle (Forsström et al., 2019; Khan et al., 2017). However, we did not observe changes in other genes classically linked to ISR and mitochondrial stress, which have been previously reported to be induced by non-exercise stressors (Forsström *et al*., 2019; Khan *et al*., 2017), such as *TRIB3*, *ATF4*, and *ATF5* mRNA (Figure S3B). We also did not observe any difference in the protein content of OMA1 (Figure S3D), which was reported to be degraded following *in vitro* mitochondrial stress (Fessler *et al*., 2020). This difference may be due to the type (i.e., exercise), magnitude (i.e., moderate number of damaged mitochondria), and duration of the stressor - SIE required as little as 3 minutes of exercise, as compared to 6-24 hours (i.e., *in vitro* experiments (Bao et al., 2016; Quiros *et al*., 2017)) or years (i.e., in mitochondrial myopathies; (Forsström *et al*., 2019; Khan *et al*., 2017)) of stress in previous studies. While further research is required, our results point to a specific ISR and mitochondrial stress response to very high-intensity exercise.

**Figure 4.**
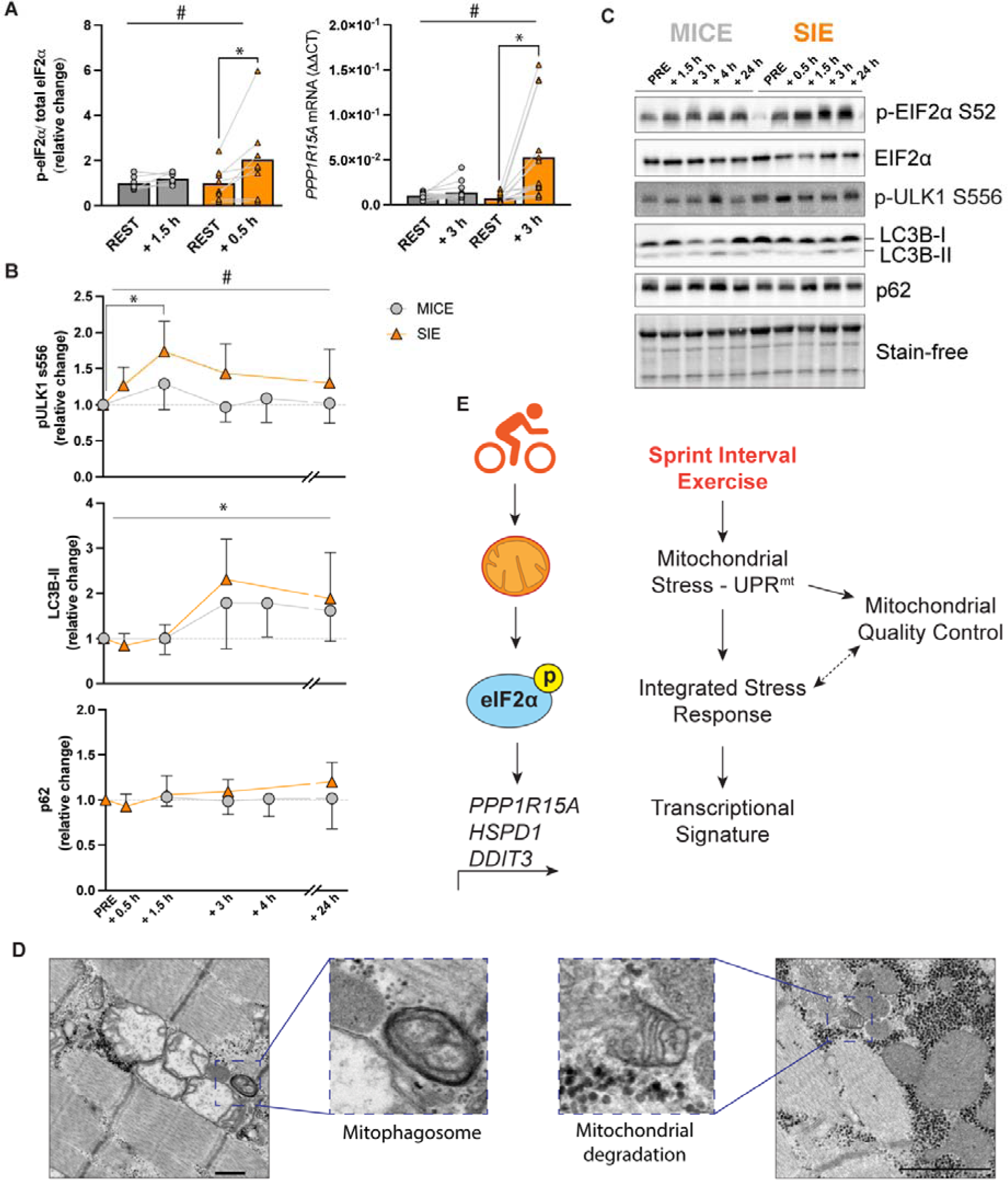
Molecular mechanisms regulating the divergent response to SIE. A) Exercise-induced increases in markers of the integrated stress response (ISR) (eIF2α phosphorylation at serine 52 [MICE n = 7; SIE n = 8]; and *PPP1R15A* mRNA [MICE n = 13; SIE n = 14] (Pakos-Zebrucka *et al*., 2016)). B) Time course of protein changes following moderate-intensity continuous exercise (MICE; n = 13 for pULK1 S556 and LC3B; n = 9 for p62) and sprint-interval exercise (SIE; n = 14 for pULK1 S556 and LC3B; n = 11 for p62) in key autophagy and mitophagy proteins. C) Representative western blots from the proteins analyzed. Stain-free image shows consistent protein loading across samples. D) Post-SIE micrograph showing swollen mitochondrial aggregates clustered together with an adjacent mitophagosome, and a mitochondrial degradation event. Scale bar = 0.5 µm. E) Model for how SIE initiates a unique transcriptional signature through the sensing of mitochondrial stress by the ISR. * denotes group differences or main effect at p < 0.05; # denotes time x group interaction at p < 0.05. For A) data are expressed as mean and for B) as mean ± 95% confidence interval. For A) and B) differences between groups were assessed via two-way ANOVA with Sidak’s multiple comparison test.

The mitochondrial ultrastructural disruption observed following SIE (Fig 3C) would suggest a need for an increased mitochondrial quality control (MQC) through mitophagy following this type of exercise. To assess mitophagy, we quantified the phosphorylation level of Unc-51-like autophagy-activating kinase 1 (ULK1) at serine 556 (due its mechanistic involvement in exercise-induced mitophagy (Laker *et al*., 2017)). Upregulated phosphorylation of ULK1 at S566 was only observed +1.5 h following SIE and displayed a group effect over the recovery period (Figure 4B and 4C). However, we did not observe any between-group differences in LC3B-II or p62 protein abundance (Figure 4B and 4C), suggesting an effect independent of general autophagy (Botella et al., 2023b). Next, we screened a recent human phospho-proteomic dataset of MICE and SIE to identify novel phosphorylation sites involved in MQC following exercise (Blazev *et al*., 2022). This dataset revealed numerous phosphorylation sites that were exclusively regulated by SIE - including eIF2α, the mitophagy receptor Optineurin (OPTN) at serine 177 (previously linked to mitophagy; (Heo et al., 2015; Lazarou et al., 2015; Richter et al., 2016; Wild et al., 2011; Wong and Holzbaur, 2014)), the autophagy receptor p62 at tyrosine 269 (associated with autophagy regulation (Linares et al., 2015)), and numerous HSPs (Figure S3C); these findings further support a larger upregulation of MQC following SIE. Using TEM, we observed multiple MQC-associated events, like mitophagosomes, mitochondrial degradation (Figure 4D), and autophagosome membrane docking on the outer mitochondrial membrane (Figure S3E), only following SIE. Collectively, these findings support our hypothesis that SIE is a potent inducer of mitochondrial stress, leading to an upregulation of the UPR^mt^ that is concurrent to the activation of the ISR and MQC pathways (Figure 4E).

### Mitochondrial biogenesis and dynamics are similarly regulated following MICE and SIE

As exercise is well-known to increase the expression of genes associated with mitochondrial biogenesis (Bishop *et al*., 2019a; Hood *et al*., 2011), and a greater exercise-induced increase in these genes remains one of the proposed mechanisms underlying the efficiency of SIE (Granata *et al*., 2017; Place *et al*., 2015), we next explored the gene expression levels of markers of mitochondrial biogenesis. We observed that the so-called master regulator of mitochondrial biogenesis, PGC1α (encoded by the *PPARGC1A* gene), was similarly increased at the transcriptional level following both types of exercise with minor between-group differences favouring either group depending on the timepoints selected (Figure S4A). There were no between-group differences in the transcriptional response of peroxisome proliferator-activated receptor alpha (PPARα) or beta (PPARβ) (Figure S4A). These results collectively suggested a largely shared transcriptional activation of common markers of mitochondrial biogenesis across MICE and SIE.

The involvement of mitochondrial dynamics components was also explored, given their proposed regulation following exercise (Huertas *et al*., 2019; Lavorato *et al*., 2018). Using a similar approach to previous research (Picard *et al*., 2013), we quantified mitochondrial contact sites from TEM micrographs (as a proxy of mitochondrial pre-fusion events) and mitochondrial pinching events (as a proxy of pre-fission events). We observed an exercise-induced effect in both mitochondrial contact sites and pinching events without between-group differences (Figure S4B). However, following SIE, we observed that the pinching events occurred specifically near the tips of mitochondria with disrupted ultrastructure (Figure S4C) - a mitochondrial fission signature that predicts mitochondrial degradation (Kleele et al., 2021). At the mRNA level, we did not observe any difference between groups in the expression of most mitochondrial dynamics factors except for mitochondrial elongation factor 2 (*MIEF2*, also known as MiD49), the gene encoding for a mitochondrial fission receptor for Dynamim-related protein 1 (DRP1; the GTPase driving mitochondrial fission). MIEF2 was divergently regulated throughout the recovery in SIE when compared to MICE, which potentially suggests an increased demand for fission (Figure S4A). However, at the protein level, we did not observe any differences for mitofusin 2 (MFN2) or fission factor 1 (FIS1) (Figure S4D). These results suggest that markers of mitochondrial biogenesis and dynamics were similarly regulated following MICE and SIE.

### There is divergent mitochondrial remodelling following moderate-intensity continuous training (MICT) and sprint-interval training (SIT)

Given the different responses observed following a single session of SIE or MICE, we next explored if these differences led to a divergent mitochondrial remodelling following training (i.e., repeated exercise sessions over weeks). To study the training-induced mitochondrial remodelling, 24 of our 28 participants completed 3 to 4 weekly exercise sessions over 8 weeks (for a total of 29 exercise sessions), and a final muscle sample was obtained 72 hours following the last exercise session (Figure 5A). There was a significant main effect of training for cardiorespiratory and endurance performance markers, such as maximal aerobic power, 4-km time-trial (TT) performance, and 20-km TT performance (all p < 0.05) (Figure S5A).

**Figure 5.**
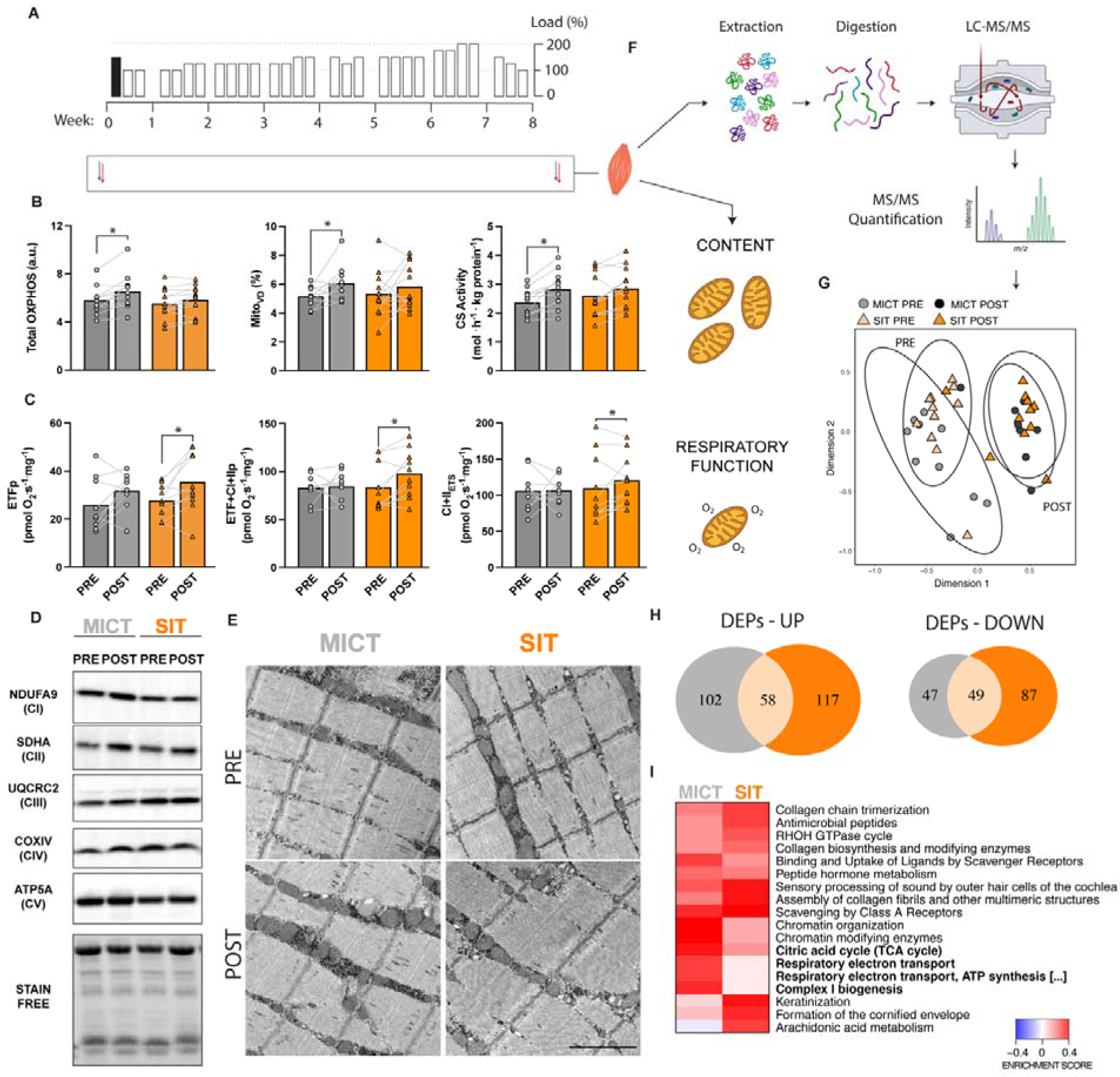
Skeletal muscle proteome remodelling differs following MICT and SIT. A) Schematic of the exercise training sessions completed throughout the 8 weeks. In black, the experimental exercise session where muscle biopsies were collected. B) Markers of mitochondrial content from whole muscle, including total oxidative phosphorylation (OXPHOS) protein content (left graph; MICT, n = 11; SIT, n = 12); mitochondrial volume density (Mito_VD_; middle graph; MICT, n = 10; SIT, n = 12), and citrate synthase activity (CS; right graph; MICT, n = 11; SIT, n = 12). C) Mitochondrial respiratory function was measured in permeabilized fibers. Left graph: respiration from electron-transferring flavoprotein (ETFp; in the presence of 0.2 mM octanyolcarnitine, 2 mM malate, 3 mM MgCl_2_ and 5 mM ADP); middle graph: respiration after addition of complex I- and complex II-linked substrates (ETF+CI+CII_P_; in the presence of 5mM pyruvate, 10 mM succinate, 3 mM MgCl2 and 5 mM ADP); right graph: electron transport system (ETS) maximal capacity obtained following titration with the uncoupler FCCP [0.7-1.5 mM]). See Material and Methods for details. D) Representative blots of proteins used to calculate total OXPHOS abundance. E) Representative micrographs from PRE and POST exercise training for each group. Scale bar = 1 µm. F) Proteomic analysis workflow. G) Principal component analysis (PCA) from the proteome analysis across both groups (MICT, n = 11; SIT, n = 12). H) Venn diagram showing that only a fraction of the differentially expressed proteins are shared across groups despite a largely similar PCA plot. I) Protein enrichment analysis across groups with using Reactome pathways. * denotes main effect at p < 0.05. Data are expressed as mean. For B) and C) data was analyzed using pre-planned one-way ANOVA.

In agreement with our previous work suggesting a divergent mitochondrial remodelling driven by either exercise volume or intensity (Bishop *et al*., 2019b; Granata *et al*., 2018; Granata *et al*., 2016a; Reisman et al., 2024), we observed that MICT led to significant increases in the overall protein content of representative subunits of the five oxidative phosphorylation complexes (OXPHOS; Figure 5B and 5D), mitochondrial volume density (Mito_VD_; Figure 5B and 5E), and citrate synthase activity (CS; Figure 5B). However, none of these markers of mitochondrial content were significantly changed following SIT (all *p* > 0.05). Of interest, the 5-fold greater training volume completed by MICT did not translate into a 5-fold greater increases in any of the markers of mitochondrial content, suggesting that the interaction of training volume with other factors, such as intensity, may be critical to maximize training-induced changes in mitochondrial content. This hypothesis is reinforced by a previous observation that no changes in Mito_VD_ were observed after 32 days of very high-volume (342 ± 42 min/day) but low-intensity cross-country skiing (Helge et al., 2008).

We next assessed the mitochondrial function, stimulated by different respiratory substrates, in permeabilized whole-muscle fibers. Following SIT, but not MICT, we observed a significant increase in the respiratory chain function from fatty acid oxidation in phosphorylating conditions (electron-transferring flavoprotein [ETFp]). Similarly, increased respiratory function was observed following the addition of complex I- and complex II-linked substrates ([ETF+CIp and ETF+CI+CII_P_], see Methods) only following SIT (*p* < 0.05; Figure 5C and S5B). This is in agreement with previous studies suggesting that exercise intensity is a key driver of changes in mitochondrial respiratory chain function (Granata *et al*., 2017). In contrast, spectrophotometric measurements of individual enzyme activities of the respiratory chain revealed that following MICT, but not SIT, there was a significant increase in Complex I (CI) activity (Figure S6B). Thus, while only MICT increased mitochondrial content and CI activity, only SIT - which was associated with an increase in mitochondrial stress and quality control pathways (see Figure 4E) - increased mitochondrial respiratory rates. Interestingly, the increased respiratory rates following SIT happened without significant changes in mitochondrial content or CI activity. These findings provide further evidence that changes in *ex vivo* mitochondrial respiratory function can be dissociated from changes in mitochondrial content and complex activity, as suggested in previous work (Granata *et al*., 2016a; Rowe et al., 2013).

### Divergent proteome-wide remodelling between MICT and SIT point to mitochondrial-specific remodelling

Having established that MICT and SIT result in distinct mitochondrial remodelling, we next used proteomics to identify if there were also protein pathways that were divergently remodelled at the whole-muscle level. We employed label-free proteomics (Figure 5F) on the whole muscle lysate and quantified 2478 proteins across all samples, which is in line with recent studies (Granata et al., 2021; Hostrup et al., 2022). Differential expression analysis showed that the protein abundance of ∼ 10 % of the detected proteins was significantly altered following SIT (250 proteins) and MICT (230 proteins) (Data S5). Principal component analysis (PCA) showed a clear effect of exercise training in both groups at the proteome level (Figure 5G), corroborating the well-known impact of exercise training on skeletal muscle plasticity (Adhihetty et al., 2003). Given the many proteins differentially regulated by each type of exercise (373 unique vs 107 shared; Figure 5H), we next employed a multidimensional gene set enrichment analysis (GSEA) approach to illustrate the coordinated or divergent training effects of MICT and SIT (Kaspi and Ziemann, 2020). Using Reactome, pathways that were most enriched following MICT, and significantly different to those enriched following SIT, were the Electron Transport Chain (ETC), the Tricarboxylic Acid Cycle (TCA), and Complex I biogenesis (Figure 5I).

These abovementioned results indicated divergent training-induced changes in the mitochondrial proteome and aligned with our biochemical and microscopy analyses. We then returned to our proteomic enrichment analysis but instead used the MitoCarta 3.0 pathways (Rath et al., 2021) – a highly curated ontology for mitochondrial proteins. Among pathways significantly enriched following both MICT and SIT, there were fatty acid oxidation, amino acid metabolism, mtRNA metabolism and carbohydrate metabolism – suggesting a generalized metabolic adaptation to MICT and SIT (Figure 6A). The pathways enriched following MICT included, not only the expected TCA cycle and ETC pathways, but also the solute carrier family SLC25A, the mitochondrial permeability transition pore, lipid metabolism, iron-sulfur containing proteins, and lysine metabolism (Figure 6A). Only a few pathways were exclusively enriched following SIT, which included folate and one-carbon (1C) metabolism, mt-tRNA synthetases, nicotinamide adenine dinucleotide (NAD^+^) biosynthesis and metabolism, and protein import, sorting and homeostasis. Some of the pathways upregulated following SIT, including 1C metabolism, NAD^+^ metabolism, and protein homeostasis, have been extensively shown to be dysregulated in *in vitro* and *in vivo* models of mitochondrial stress (Bao *et al*., 2016; Forsström *et al*., 2019; Khan *et al*., 2017). Therefore, our proteomic results suggest that the mitochondrial stress and UPR^mt^ observed following SIE may influence the mitochondrial proteome remodelling to exercise training. Furthermore, this divergent mitochondrial-specific proteome remodelling following MICT and SIT also aligns with the notion that specific mitochondrial remodelling may be achieved when different exercise regimes are prescribed (Granata *et al*., 2021).

**Figure 6.**
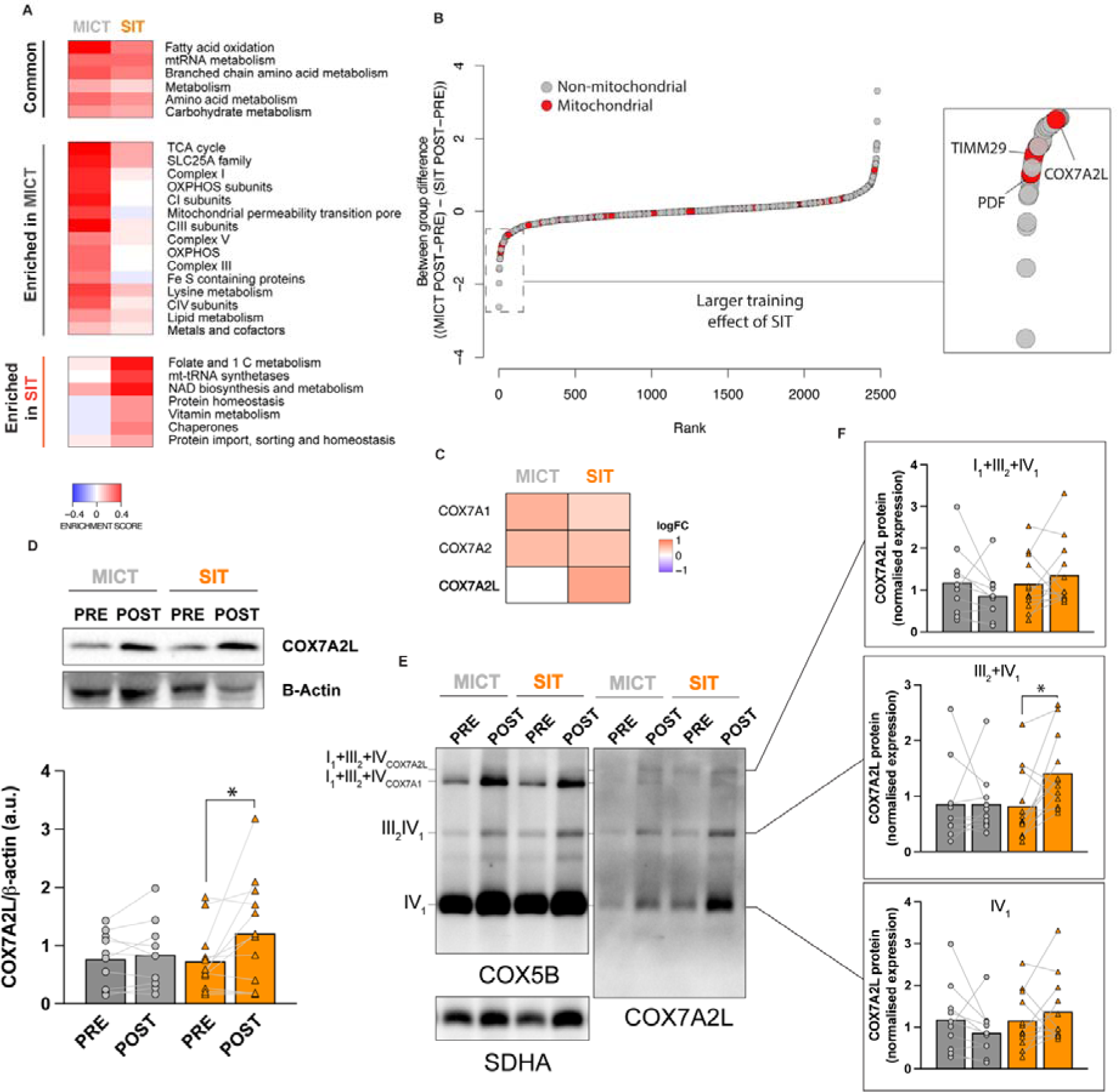
Mitochondrial-specific proteome remodelling reveals divergent supercomplex remodelling driven by COX7A2L protein. A) Two-dimension differential protein enrichment analysis using mitch package and utilizing MitoCarta 3.0 gene sets. B) Comparative analysis of training-induced protein relative abundance changes following MICT and SIT across individual proteins; proteins defined as mitochondrial from MitoCarta 3.0 are highlighted in red. C) COX7A2L was identified as a divergently regulated protein between MICT (n = 10) and SIT (n = 12), also compared to other COX7A family members. D) Representative SDS-PAGE blot of COX7A2L and the housekeeping protein ß-actin from the whole-muscle analysis. The lower graph represents COX7A2L protein levels in whole-muscle lysates PRE and POST MICT and SIT, showing a significant increase following SIT. E) Representative BN-PAGE blot of the relative COX7A2L abundance and distribution across respiratory chain complexes and supercomplexes following MICT (n = 10) and SIT (n = 12). F) Quantitative changes in the normalized (by citrate synthase activity) protein abundance of COX7A2L in each supercomplex compartment. * denotes main effect at p < 0.05. Data are expressed as mean. For D) and F) differences between groups were assessed via pre-planned one-way ANOVA.

Lastly, we wondered how SIT could lead to an increased mitochondrial respiratory chain function without a concurrent increase in markers of mitochondrial content or OXPHOS protein abundance. To this end, we took advantage of our extensive proteome-wide analysis and ranked proteins according to their between-group training-induced differences (Figure 6B and Data S6). We focused our attention on mitochondrial proteins with a greater magnitude of change following SIT when compared to MICT. Among the top three mitochondrial proteins found, peptide deformylase (PDF), translocase of inner mitochondrial membrane 29 (TIMM29), and cytochrome C oxidase subunit 7A2 like (COX7A2L; also called supercomplex assembly Factor 1, SCAF1), were identified. These proteins are involved in mitochondrial protein synthesis (Escobar-Alvarez et al., 2010), protein import through the inner membrane (Callegari et al., 2016), and assembly of respiratory chain supercomplexes (SCs) (Fernández-Vizarra and Ugalde, 2022), respectively.

We focused our attention on COX7A2L (Figure 6C), which has previously been associated with markers of muscle function (Benegiamo et al., 2022). COX7A2L was also an attractive candidate because its expression is induced following the activation of the ISR (Balsa et al., 2019) (also upregulated to a larger extent following SIE; see Figure 4). Furthermore, COX7A2L has been proposed to promote the formation of SC III_2_+IV_1_ and of the S-respirasome (SC I_1_+III_2_+IV_1-2_ containing COX7A2L) under conditions of greater glycolytic metabolism (Fernández-Vizarra et al., 2022), and our own data highlights the greater activation of glycolysis with SIE compared to MICE (Figure 1C). Finally, we did not observe a protein enrichment of any other OXPHOS components exclusively following SIT. Immunoblotting analyses confirmed the significant increase of COX7A2L protein abundance only following SIT (Figure 6D), but not MICT, and this was independent of changes in markers of mitochondrial content (Figure 5B). In contrast, the level of the canonical COX7A1 protein was significantly increased only following MICT (Figure 6C and Data S6). This observation suggested the possibility of a COX7A-mediated reorganization of the respiratory chain complexes and SCs to adapt mitochondrial respiratory function to the divergent metabolic conditions induced by MICT (more oxidative) and SIT (more glycolytic).

We are not aware of previous studies that have investigated the potential role of COX7A2L in training-induced mitochondrial adaptations in humans. Research has shown that COX7A2L is essential to drive the formation of distinct supercomplex structures (Fernández-Vizarra *et al*., 2022; Vercellino and Sazanov, 2021; Vercellino and Sazanov, 2024). Evidence from previous studies showed that the canonical COX7A1/2 proteins are exchangeable. Under conditions promoting oxidative metabolism the formation of the more efficient C-MRC organization is favoured (I_1_+III_2_+IV_1-2_ containing COX7A1/2, free CIII_2_, and CIV). Conversely, under conditions that promote glycolytic metabolism, COX7A2L plays a key role in promoting the formation of the less-efficient S-MRC (I_1_+III_2_+IV_1-2_ containing COX7A2L and SC III_2_+IV_1_) (Fernández-Vizarra *et al*., 2022). Using blue-native PAGE (Figure 6E and S6A), we explored if MICT- or SIT-induced changes in specific COX7A isoforms were associated with diverse changes in SCs and respirasomes. Notably, we observed that the SIT-induced increase in COX7A2L levels was associated with a significant accumulation of SC III_2_+IV_1_ but not of the COX7A2L-containing respirasomes (I_1_+III_2_+IV_1-2_; Figure 6F). It is possible that this greater accumulation of SC III_2_+IV_1_ only following SIT contributed to the significant increase in mitochondrial respiratory function in response to SIT but not MICT (Figure 5C), supported by previous structural studies showing that CIII_2_ and CIV gain catalytic advantage when assembled into this supercomplex (Vercellino and Sazanov, 2021). However, there were no significant correlations between relative changes in COX7A2L protein levels (at the whole muscle level and within each SC) and changes in mitochondrial respiratory function following SIT (Figure S6C), suggesting that other factors may be contributing to the enhanced respiratory function observed following SIT.

## DISCUSSION

Here, we show that sprint-interval exercise (SIE) uniquely provokes mitochondrial ultrastructural disturbances that precede a transcriptional signature of mitochondrial stress and the activation of the mitochondrial unfolded protein response (UPR^mt^). These changes, observed only following SIE and not MICE, were also associated with a greater activation of the integrated stress response (ISR) and mitochondrial quality control (MQC) pathways. These distinct acute responses were complemented by divergent mitochondrial remodelling - moderate-intensity continuous training (MICT) was characterized by an increase in markers of mitochondrial content while sprint interval training (SIT) leads to increased mitochondrial respiratory chain function. Subsequent proteomic analyses revealed a unique enrichment of pathways involved in One-Carbon (1C), NAD^+^ metabolism, and mitochondrial protein quality control pathways, following SIT. We also identified a COX7A2L-mediated accumulation in the III_2_+IV_1_ respiratory chain supercomplex exclusively following SIT. Our results provide valuable new insights on the molecular mechanisms by which distinct mitochondrial remodelling can be achieved using different exercise prescriptions. Our findings support a novel concept whereby very high-intensity sprint exercise provokes mitochondrial structural disturbances and activates mitochondrial-specific stress responses in human skeletal muscle.

The health benefits of exercise are well known despite the underlying molecular mechanisms remaining partially unresolved (Zierath and Wallberg-Henriksson, 2015). It is widely understood that the exercise prescription (e.g., the choice of exercise intensity) is a crucial variable to optimize the benefits of exercise (Haskell et al., 2007; Lee et al., 1995). Here, we show that both SIE and MICE elicit a mostly shared transcriptional response to exercise (Figure 1). However, SIE produced a unique molecular signature early in the recovery, which was characterized by the upregulation of the UPR^mt^ and related mitochondrial stress pathways (Figure 2). While this aligns with research in mice reporting that exercise activates the endoplasmic reticulum unfolded protein response (UPR^ER^) (Wu et al., 2011), the UPR^mt^ has remained unexplored in most exercise studies – especially those involving humans. Our results suggest that the mitochondrial disturbance following SIE, but not MICE, is characterized by mitochondrial ultrastructural and morphological disturbances that likely contributes to the activation of the UPR^mt^, as well as the ISR and MQC pathways. The adaptive nature of the UPR^mt^ seems to be largely dependent on the tissue, persistence, and reversibility of the stressor (Sprenger and Langer, 2019). In fact, skeletal muscle myopathies involving mitochondrial genetic defects are characterized by a chronic mitochondrial stress that is seemingly maladaptive (Forsström *et al*., 2019; Khan *et al*., 2017). In contrast, we hypothesize that the short and transient nature of the stress invoked by sprint interval exercise makes it a potent mitohormetic event leading to adaptive skeletal muscle remodelling. The transient nature of the mitochondrial stress is supported by our observation that the mitochondrial stress signature elicited by SIE was no longer present 24 hours following exercise (Figure S2). Our results may also help explain previous reports suggesting a mitochondrial functional impairment in skeletal muscle when biopsies are taken shortly after periods of excessive high-intensity training (Cardinale et al., 2021; Flockhart et al., 2021) but not when sufficient recovery is provided prior to the muscle sampling (Granata *et al*., 2021; Granata et al., 2016b). It is tempting to suggest that repeated mitochondrial stress, without sufficient recovery between sessions, could potentially underlie the maladaptive nature of certain exercise training programs and should be queried in future studies. Given the large body of evidence suggesting that high-intensity interval training is more effective at improving markers associated with a wide variety of metabolic and age-related diseases, it remains to be fully elucidated whether the activation of UPR^mt^, which appears largely specific to sprint-interval exercise in the present study, may also underlie some of these benefits.

Increased mitochondrial content and respiratory chain function following endurance training have long been known and are often depicted as stoichiometric adaptations (Holloszy, 1967). However, contrary to this long-held view, our results indicate that different exercise prescriptions produce specific and divergent alterations to mitochondrial content, respiratory chain function, and the mitochondrial proteome (Figure 5). MICT led to increases in multiple markers of mitochondrial content and CI enzymatic activity, none of which were significant following SIT, in agreement with previous literature (Granata *et al*., 2017; Meinild Lundby et al., 2018). In contrast, SIT was characterized by a functional increase in mitochondrial respiratory chain function and a specific remodelling of the mitochondrial proteome that was linked to pathways related to NAD^+^ metabolism, protein import, protein folding, and 1C metabolism (Figure 5); these are all crucial pathways in the activation and response to the UPR^mt^ (Bao *et al*., 2016; Sutandy *et al*., 2023), which was also largely specific to sprint-interval exercise in the present study. Whether the mitochondrial structural disturbances, and the increased activation of mitochondrial stress and MQC pathways, may have contributed to the improvement in mitochondrial respiratory function through enhanced mitophagy (e.g., selective removal of ‘less fit’ mitochondria) warrants further research.

Another unique adaptation to SIT was a significant increase in COX7A2L protein abundance, which accumulated primarily in the III_2_+IV_1_ supercomplex (Figure 6). This is consistent with research demonstrating that COX7A2L-mediated accumulation of III_2_+IV_1_ supercomplexes is a preferential structural organization in cells relying primarily on glycolytic metabolism (Fernández-Vizarra *et al*., 2022), as occurred following SIE (see Figure 1C). Furthermore, previous research has linked COX7A2L-mediated supercomplex assembly in mammalian cells to the activation of the ISR (Balsa *et al*., 2019), which was upregulated to a larger extent following SIE in the present study (Figure 4). Interestingly, ISR activation has been shown to diminish the levels of TCA cycle intermediate metabolites while leading to an accumulation of glycolytic intermediates (Labbé et al., 2024). Thus, we speculate that the COX7A2L-mediated remodelling observed following SIT may help sustain mitochondrial bioenergetics under stress conditions via facilitating glycolytic flux.

A final observation is that the greater accumulation of SC III_2_+IV_1_ following SIT was not associated with the greater increase in respiratory chain function following SIT (Figure S6C). While the functional relevance of SCs continues to be debated, our findings are consistent with suggestions that the main role of SC formation is to provide structural stability rather than to enhance electron transfer between complexes during respiration (Brischigliaro et al., 2023; Fernández-Vizarra *et al*., 2022; Milenkovic et al., 2023). Thus, while an increase in the abundance of COX7A2L and COX7A2L-containing SC III_2_+IV_1_ was specific to SIT, the functional implications require further investigation. More research is required to resolve whether the supramolecular regulation of mitochondrial complexes influences mitochondrial bioenergetics following exercise training in humans (Granata *et al*., 2021; Greggio et al., 2017).

In summary, our findings support the notion that sprint interval exercise is a natural mitohormetic stressor in humans. We further highlight the importance of personalized training interventions to achieve specific mitochondrial remodelling and provide insights into the molecular responses to different exercises. As many of the health benefits of exercise are believed to originate from changes in mitochondria, these results may be used to optimize the prescription of exercise for targeted mitochondrial adaptations important for human health.

### Limitations of Study

The recruitment of young healthy males in this study means that extrapolation of our results to other populations should be done with caution. Future research should explore whether the differences observed in the present study are also observed in females and older participants, under different exercise prescriptions, and in pathological conditions such as metabolic disease. Future studies should use *in vitro* and *in vivo* models of exercise to untangle the precise mechanisms by which mitochondria sense and communicate the stress to regulate the adaptive response to exercise training observed following sprint interval training.

## Methods

### Participants and ethics approval

Twenty-eight healthy males (26.5 ± 5.3 y; 179.1 ± 6.3 cm; 76.8 ± 10.3 kg) volunteered to take part in this study. Participants were informed of the study requirements, benefits, and risks involved before giving their written informed consent. Participants were matched for their maximum aerobic power (W[_max_; W.kg^−1^) and assigned in a random, counter-balanced order to one of the two exercise groups. All participants completed a single experimental exercise session. Two participants from the MICT group and two from the SIT group withdrew from the training study due to time constraints, and the data from one MICT participant was excluded from the final analysis as their final muscle sample was of poor quality. Ethics approval for the study was obtained from the Victoria University Human Research Ethics Committee (HRE17-075) and was registered as a clinical trial under Australian New Zealand Clinical Trials Registry (ANZCTR; ACTRN12617001105336).

### Graded exercise test (GXT)

The GXTs were conducted using an electronically braked cycle ergometer (Lode Excalibur v2.0, The Netherlands). A metabolic analyser (Quark Cardiopulmonary Exercise Testing, Cosmed, Italy) was used during the testing to assess V[O_2_, V[CO_2_, V[_E_, on a breath-by-breath basis, and heart rate was also measured (Garmin, USA). A GXT with 1-min stages was chosen (Jamnick et al., 2018), and the increase in power between stages was adjusted following the familiarisation trial to attain a total duration of 9 to 11 min upon exhaustion. After reaching exhaustion, 5 min of rest were provided before a verification bout (time to exhaustion at 90 % of W[_max_) was applied to verify that V[O_2max_ was achieved.

### 4-km Time Trial (TT) and 20-km Time Trial (TT)

Time trials were used as a marker of endurance performance and were completed on an electronically-braked cycle ergometer (Velotron, RacerMate, Seattle, WA, USA). Prior to each time trial, participants completed a 5-min warm-up at a self-selected intensity followed by 5 min of rest. During the time trial, participants were only allowed to see the bike gearing and speed, but not completed or remaining distance, and were constantly provided with encouragement throughout the trial. Performance data were expressed as absolute mean power (W), and relative mean power (W.kg^−1^)

### Submaximal test

Following the completion of the GXT, the ventilatory parameters obtained (V[O_2_, V[CO_2_, V[_E_) were plotted and visually inspected to estimate the first ventilatory threshold (VT1). Accordingly, a power 40 W lower than the estimated VT1 was selected as the starting point of the submaximal test; 3-min stages, with the intensity increased by 10 W every 3 min until the first lactate threshold (LT1) could be identified. LT1 was defined as the power associated with the first increase in lactate that was at least 0.3 mmol.L^−1^ greater than the previous stage during the submaximal test. Antecubital venous blood was taken in the last 15 s of each stage and was analyzed using a blood lactate analyzer (YSI 2300 STAT Plus, YSI, USA).

### Sprint-Interval Exercise (SIE) or Training (SIT)

Fourteen participants were randomly allocated to this group (26.9 ± 4.9 y; 179.6 ± 5.9 cm; 76.5 ± 9.7 kg; 23.7 ± 2.8 BMI). The exercise session consisted of six, 30-s, ‘all-out’ cycling bouts against a resistance initially set at 0.075 kg.kg BM^−1^, interspersed with a 4-min recovery period (Granata *et al*., 2016a). During the recovery participants remained on the bikes and were allowed to either rest or cycle against no resistance. During the last 30 s of the recovery period participants were instructed to begin pedalling and to reach a cadence of 90 rpm against no resistance, and in the last 10 seconds they were advised to get ready and the countdown began. They were instructed to begin pedalling as fast and hard as possible when the countdown reached zero. At this time, the load was applied via the ergometer software (Lode Excalibur v2.0, The Netherlands). Participants were verbally encouraged to pedal as fast and hard as possible during the entire duration of the bout. For the training period, participants progressively completed up to 8 sprints per session in week 7. The resistance load was increased to 0.080 kg.kg BM^−1^ in week 3, to 0.085 kg.kg BM^−1^ in week 5, and to 0.090 kg.kg BM^−1^ in week 7.

### Moderate-Intensity Continuous Exercise (MICE) or Training (MICT)

Fourteen participants were randomly allocated to this group (26.1 ± 5.7 y; 178.6 ± 6.5 cm; 77.0 ± 10.7 kg; 24.1 ± 2.6 BMI). The exercise session consisted of continuous cycling at a fixed power equivalent to ∼ 90 to 100 % of LT1. The duration of the experimental session was 90 min. During the training period, participants progressively completed up to 120 min per session in week 7, and the intensity was reassessed and adjusted accordingly (using the submaximal test) every 2 weeks.

#### Familiarisation and testing period

Following the initial graded exercise test (GXT), participants were allocated to their exercise group. During the following two weeks, participants completed multiple familiarisation sessions including a 4-km time trial (4km-TT), a 20-km time trial (20km-TT), a submaximal test, and the exercise session of their respective group twice. Following this, participants underwent a week of testing (GXT, 4km-TT and 20km-TT). Participants were required to avoid any vigorous exercise for at least 48 h preceding each performance test, to avoid caffeine consumption for at least 8 h prior to each test, and to continue their habitual dietary pattern. Tests were performed at a similar time of the day throughout the study to avoid any influence of circadian rhythms.

### Single exercise session trial

#### Nutritional standardisation

Participants were requested to maintain their habitual dietary pattern and physical activity throughout the study. To minimize between-subject variation, participants were provided with standardized caloric and macronutrient intake for the 24 h leading to the experimental exercise trial. Participants were provided with a standardized dinner (55 kJ/kg of body mass (BM), providing 2.1 g carbohydrate/kg BM, 0.3 fat/kg BM, and 0.6 g protein/kg BM) and breakfast (41 kJ/kg BM, providing 1.8 g carbohydrate/kg BM, 0.2 g fat/kg BM, and 0.3 g protein/kg BM) to be consumed 15 h and 3 h before the biopsies, respectively. These two standardized meals were provided before the experimental exercise trial, as well as before the + 24 h biopsy. Participants were also asked to refrain from caffeine during the day of the trial.

#### Muscle biopsies

All muscle samples were obtained by an experienced medical doctor at a similar time of the day (morning), at a constant depth of around 2 to 3 cm, and under local anaesthetic injected into the skin and fascia (1% xylocaine, Astra Zeneca). Muscle biopsies were taken from the *vastus lateralis* muscle using the Bergström biopsy needle technique with suction. Once the muscle sample was obtained it was processed, cleaned of excess blood, fat, and connective tissue, and split in multiple portions. One portion (5 to 10 mg) was immediately fixed in a 0.2 M Na-Cacodylate buffered 2.5% glutaraldehyde and 2% paraformaldehyde for transmission electron microscopy imaging. One portion (10 mg) was immediately immersed in a tube containing biopsy preserving solution (BIOPS) kept on ice, and then used for *ex vivo* measurements of mitochondrial respiratory function. The remaining portion was immediately frozen in liquid nitrogen and stored at −80 °C for subsequent analyses.

### Biochemical analyses

#### Blood Lactate and pH measurements

Antecubital venous blood samples (∼ 1 mL) were collected pre and post the first exercise session from a cannula inserted in the antecubital vein for the determination of venous blood hydrogen concentrations [H+] and lactate concentrations using a blood-gas analyzer (ABL 800 FLEX, Radiometer Copenhagen).

##### Western Blotting

Approximately 10 to 20 mg of frozen muscle was homogenized 2 times for 2 minutes at a speed of 30 Hz with a TyssueLyser instrument (Qiagen, Canada) in an ice-cold lysis buffer (1:20) containing 50 mM Tris-HCl, 150 mM NaCl, 1 mM EDTA, 5 mM Na4P2O7, 1 mM Na3VO4, 1% NP-40, with added protease and phosphatase inhibitors at a 1:100 concentration (Cell Signaling Technology). Protein concentration was determined using a commercial colorimetric assay (Bio-Rad Protein Assay kit II, Bio-Rad Laboratories Pty Ltd, Gladesville, NSW, AUS) and lysates were then diluted with an equal volume in 2x Laemmli buffer containing 10% β-mercaptoethanol. For each protein of interest a linearity study was conducted to determine the ideal loading amount. Muscle lysates were then loaded in equal amounts (10 to 20 μg according to target protein) and separated by electrophoresis for 1.5 to 2.5 h at 100 V using pre-cast SDS-PAGE gels (4-20%). Once resolved, the gels were then wet transferred onto PVDF LF or nitrocellulose membranes using a Turbo Transfer (Bio-rad Laboratories Pty Ltd, Gladesville, NSW, AUS). When appropriate, membranes were imaged for total protein, which was used later as a loading control. Membranes were then blocked at room temperature for 1 h using 3% skim milk or 3% Bovine Serum Albumin (BSA) in Tris Buffer Saline (TBS) 0.1% Tween-20 (TBS-T). After 3 x 5-min washes in TBS-T, membranes were incubated overnight at 4°C with gentle agitation in primary antibody solutions (3% BSA plus 0.02% Na Azide). Immunoblotting was carried out using the desired antibody as follows: pULK1 S556 (CST# 5869), LC3B (CST #3868S), P62 (#ab56416), OMA1 (#sc-515788), FIS1 (#sc-376469), MFN2 (CST #9482), UQCRC2 (#ab14745), COX IV (#ab14744), SDHA (#ab14715), NDUFA9 (#ab14713), ATP5A (#ab14748), EIF2a (CST #5324), pEIF2a S52 (CST #3398), COX7A2L (ProteinTech #11416-1-AP), B-Actin (Sigma #A1978), COX5B (#sc-374417). The following morning, membranes were washed 3 x 5 min in TBS-T and subsequently incubated under gentle agitation at room temperature with the appropriate secondary antibody for 60 to 90 min in 5% skim milk in TBS-T. Membranes were then washed again for 3 x 5 min in TBS-T before being immersed for 5 min under gentle agitation at room temperature in Clarity ECL detection substrate (Bio-rad Laboratories Pty Ltd, Gladesville, NSW, AUS). Protein bands were visualized using a Bio-Rad ChemiDoc imaging system and band densities were determined using Bio-Rad Image Lab analysis software (Bio-Rad Laboratories Pty Ltd, Gladesville, NSW, AUS). Finally, all samples for each participant were loaded on the same gel and the different concentrations of a mixed-homogenate internal standard (IS) were also loaded on each gel and a calibration curve plotted of density against protein amount. From the subsequent linear regression equation, protein abundance was calculated from the measured band intensity for each lane on the gel. Total protein content of each lane was obtained from the stain-free image of the membrane and was used for normalization of the results.

#### Blue Native Electrophoresis and In-Gel Activity Assay (IGA)

Mitochondrial pellets were isolated from enriched fractions obtained from skeletal muscle homogenates. Briefly, ∼ 20 mg of the muscle biopsy were cut in small pieces using a surgical scalpel. The tissue was then homogenized in a glass-glass Dounce type potter using 20 volumes of Medium A (0.32 M sucrose, 10 mM Tris-HCl pH=7.4, 1 mM EDTA) and 15 manual strokes. The homogenate was transferred to an Eppendorf tube and centrifuged at 800 g during 5 minutes at 4 °C. The pellet was discarded, and the supernatant was collected. Half of the supernatant was transferred to a clean tube and kept for the determination of the enzymatic activities (see below). The other half was transferred to a different tube and centrifuged at 9000 g for 10 minutes at 4 °C. The supernatant was discarded and the pellet containing the isolated mitochondria was solubilized using digitonin at a detergent-to-protein ratio of 4:1. Pre-cast Native PAGE 3-12% Bis-Tris gels (Invitrogen) were loaded with 60-80 µg of mitochondrial protein and processed for blue native electrophoresis (BN-PAGE), as previously described (Timón-Gómez et al., 2020). After electrophoresis, proteins were transferred to PVDF membranes at 40 V overnight and probed with antibodies. Duplicate gels were used for CI-IGA assays.

#### Reverse transcription and quantitative polymerase chain reaction (qPCR)

For each sample, 1 μg of RNA was transcribed into cDNA on a thermal cycler (S1000TM Thermal Cycler, Bio-Rad, USA) using the iScriptTM cDNA Synthesis Kit (Bio-Rad, USA) and the following incubation profile: 5 min at 25 °C, 30 min at 42 °C and 5 min at 85 °C. The transcription was performed with random hexamers and oligo dTs in accordance with the manufacturer’s instructions. A reverse transcriptase (RT) negative control was also generated.

Forward and reverse primers for the target and housekeeping genes were designed based on NCBI RefSeq using NCBI Primer-BLAST (www.ncbi.nlm.nih.gov/BLAST/) or obtained from scientific publications. Specificity of the amplified product was confirmed by melting point dissociation curves. The mRNA expression was performed by quantitative real-time RT-PCR (Mastercycler® RealPlex2, Eppendorf, Germany), using a 5 μL PCR reaction volume with SYBR Green supermix (SsoAdvanced™ Universal SYBR® Green Supermix, Bio-Rad, USA). All samples were run in duplicate simultaneously with template free controls, using an automated pipetting system (epMotion 5070, Eppendorf, Germany). The following PCR cycling patterns were used: initial denaturation at 95 °C (3 min), 40 cycles of 95 °C (15 s) and 60 °C (60 s). The mRNA expression of five housekeeping genes was quantified, and the three most stable genes were determined using the BestKeeper software (Pfaffl et al., 2004). 18S ribosomal RNA (18s), glyceraldehyde 3-phosphate dehydrogenase (GAPDH), and beta-2-microglobulin (B2M) were classified as most stable and utilized for the analysis. Expression of each target gene was calculated as previously published (Kuang et al., 2018).

##### Mitochondrial respiratory chain complex activity

Half of the supernatant obtained after the 800xg centrifugation of the skeletal muscle homogenates (see above—Blue Native Electrophoresis sample preparation) was transferred to a clean tube and immediately frozen at −80 °C and kept overnight. Before performing the measurements, the samples were thawed at 37 °C and snap-frozen again in liquid nitrogen, repeating the freeze-thawing cycle one more time (three cycles in total). The measurements of the rotenone-sensitive NADH dehydrogenase activity (complex I) and of the cytochrome c oxidase activity (complex IV) were performed using 10-30 µL of mitochondria-enriched supernatant, in a total reaction volume of 200 µL, using a plate-reader spectrophotometer as described (Brischigliaro et al., 2022).

##### Citrate Synthase (CS) Activity Assay

Using the whole-muscle lysates, CS activity was determined in triplicate on a microtiter plate by adding: 5 μL of a 2 mg.mL^−1^ muscle homogenate, 40 μL of 3 mM acetyl CoA in Tris buffer and 25 μL of 1 mM 5,5’-dithiobis(2-nitrobenzoic acid) (DTNB) in Tris buffer to 165 μL of 100 mM Tris buffer (pH 8.3) kept at 30°C. At this point 15 μL of 10 mM oxaloacetic acid were added to the cocktail and the plate was immediately placed in a spectrophotometer kept at 30°C (xMark Microplate Spectrophotometer, Bio-Rad Laboratories Pty Ltd, Gladesville, NSW, AUS). Following 30 s of linear agitation, absorbance at 412 nm was recorded every 15 s for 3 min. CS activity was calculated and reported as mol.kg protein^−1.^h^−1^.

#### Preparation of permeabilised skeletal muscle fibers

A 10 mg fresh muscle sample was placed in ice-cold BIOPS, a biopsy preserving solution containing 2.77 mM CaK2EGTA, 7.23 mM K2EGTA, 5.77 mM Na2ATP, mM 6.56 MgCl2, 20 mM taurine, 50 mM MES, 15 mM Na2 phosphocreatine, 20 mM imidazole, and 0.5 mM DTT adjusted to pH 7.1. Samples were mechanically separated using pointed forceps while kept on ice. Fibers were subsequently permeabilized by gentle agitation for 30 min at 4 °C in BIOPS containing 50 μg.mL^−1^ of saponin. Samples were then washed for 3 x 7 min at 4 °C by gentle agitation in MiR05, a respiration medium containing 0.5 mM EGTA, 3 mM MgCl2, 60 mM K-lactobionate, 20 mM taurine, 10 mM KH2PO4, 20 mM Hepes, 110 mM sucrose and 1 g.L^−1^ BSA essentially fatty acid-free adjusted to pH 7.1 at 37 °C. This method selectively permeabilizes the cellular membrane leaving the mitochondria intact and allows for *ex vivo* measurements of mitochondrial respiration.

#### Mitochondrial respiratory function

After washing, 2 to 4 mg wet weight of muscle fibers were assayed in triplicate in a high-resolution respirometer (Oxygraph-2k, Oroboros Instruments, Innsbruck, Austria) containing 2 mL of MiR05. Mitochondrial respiration was measured at 37 °C. Oxygen concentration (nmol.mL^−1^) and oxygen flux (pmol.s^−1.^mg^−1^) were recorded using DatLab software (Oroboros Instruments, Innsbruck, Austria), and instrumental background oxygen flux, accounting for sensor oxygen consumption and oxygen diffusion between the medium and the chamber boundaries, was corrected online. Re-oxygenation by direct syringe injection of O_2_ in the chamber was necessary to maintain O_2_ levels between 275 and 450 nmol.mL^−1^, so as to avoid a potential oxygen diffusion limitation. The following substrates and inhibitors were added as following: octanyolcarnitine (0.2 mM) and malate (2 mM) were added to measure the LEAK respiration from electron-transferring flavoprotein (ETF_L_); this was followed by addition of MgCl2 (3 mM) and ADP (5 mM) for the measurement of phosphorylation capacity (P) through ETF (ETF_P_); the subsequent addition of pyruvate (5 mM) allowed the measurement of complex I and ETF respiration (ETF+CI_P_); this was followed by addition of succinate (10 mM), which provided the measurement of P through complex I and complex II (ETF+CI+CII_P_); after these steps cytochrome c (10 mM) was added to test for the outer membrane integrity, followed by a stepwise carbonyl cyanide 4-phenylhydrazone (FCCP) titrations (0.7-1.5 mM) to obtain the electron transport system capacity (E) through CI+CII (CI+CII_ETS_). Antimycin A (2.5 mM), an inhibitor of complex III, was then added for the measurement and correction of residual O_2_ consumption as a measure of non-mitochondrial O_2_ consumption.

#### Transmission Electron Microscopy

Skeletal muscle samples were fixed overnight at 4 °C with 0.2 M sodium cacodylate– buffered, 2.5% glutaraldehyde, and 2% paraformaldehyde. Fixed samples were rinsed with 0.1 M sodium cacodylate, and postfixed with ferricyanide-reduced osmium tetroxide (1 % OsO4, 1.5 % K3 [Fe(CN)6], and 0.065 M cacodylate buffer) for 2 h at 4 °C. The postfixed samples were rinsed with distilled water, and then stored overnight in 70% ethanol. Dehydration was performed by graduated ethanol series (80%, 90%, 95%, 100%, and 100%; 10 min each) and propylene oxide (100% and 100%; 5 min each). Samples were infiltrated with Araldite 502/Embed 812 by graduated concentration series in propylene oxide (25% for 1 h, 33% for 1 h, 50% overnight; 66% for 4 h, 75% for 4 h, 100% overnight; and 100% for 5 h) and then polymerized at 60 °C for 48 h. Embedded samples were sectioned using an Ultracut UCT ultramicrotome (Leica Biosystems) equipped with a 45 ° diamond knife (Diatome) to cut 75-nm ultrathin sections. The grids were stained at room temperature using 2% aqueous uranyl acetate (5 min) and Reynolds lead citrate (3 min) before routine imaging. All TEM imaging was performed at 80 kV on a Hitachi H-7500 TEM using a Gatan 791 MultiScan side-mount CCD camera and DigitalMicrograph (Version 1.71.38) acquisition software. All images were obtained from the longitudinal section. Image acquisition was performed by a blinded investigator.

#### Mitochondrial volume density

Forty micrographs of approximately 13 x 13 µm were acquired from at least five different muscle fibers from each sample. A total of twenty micrographs were randomly selected and quantified using the Cavalieri stereology method to estimate volume. The grid spacing used was 0.5 µm for both axes and was selected accordingly to a previously published methodological study (Broskey et al., 2013). Mitochondrial volume density was then expressed as a percentage of the grid intersections that overlapped with a mitochondrion relative to the total number of grid intersections.

#### Mitochondrial morphology and structure

To assess mitochondrial morphology, ten images were randomly acquired, from at least three different fibers, in the PRE and + 0 h samples from the subset of participants that had fixed post-exercise samples (n = 4 in MICE, n = 5 in SIE). Mitochondrial contact number was calculated as the frequency that two adjacent mitochondria had membrane contact relative to total number of mitochondria (total number of visualized mitochondria = 4923), obtained from micrographs with enough quality to detect such contacts (n = 184). Mitochondria shape and size descriptors were obtained using ImageJ (NIH, USA), as previously published (205), by manually tracing clear mitochondria (MICE PRE, n = 428; MICE + 0 h, n = 446; SIE PRE, n = 603; SIE + 0 h, n = 551). Frequency distribution plots were generated using the *ggplot* package in R. Disturbed mitochondria were evaluated by counting the mitochondria from micrographs (micrographs = 207; total number of visualized mitochondria = ∼ 5400; disturbed mitochondria = 228). Given the striking morphological and ultrastructural difference in mitochondria following SIE, whole tile-sets of at least 4 individual fibers from each post-SIE sample (n = 8) were imaged and visually inspected to confirm these exercise-induced morphological and ultrastructural changes following SIE. Mitochondrial cristae variables were obtained using ImageJ (NIH, USA). To calculate cristae density, a line was drawn spanning an entire mitochondrion (and perpendicular to cristae orientation) and the number of cristae that crossed this line (#) was calculated per mitochondrion length (µm). For cristae junctions, the area (µm^2^) of a given mitochondrion was measured and the number of cristae junctions near the outer mitochondrial membrane were counted (#). For cristae width, the average of three perpendicular measurements (membrane to membrane) from each individual crista were used to establish the cristae width.

### Bioinformatic analysis

#### RNA-Sequencing

Transcriptomic analyses were performed by the Beijing Genomic Institute (BGI). Sequencing was performed using 100bp paired-end sequencing on the DNBseq technology platform. A total of 1 μg RNA from each of the 72 samples (24 participants and 3 time-points) were used. From the raw reads obtained, quality control was performed using FastQC. Reads were then aligned against the GRCh38 Human genome assembly using STAR 2.7.3a. Reads were mapped to their corresponding genes using RSEM. The read counts obtained were then used in R for differential gene expression with limma and DESeq2 package following previously published guidelines. Geneset enrichment analysis of mitostress geneset (Quiros *et al*., 2017) was performed using the fGSEA R package (Korotkevich et al., 2021). The data will be deposited at Gene Expression Omnibus upon publication.

#### Proteomics

The skeletal muscle samples (n = 46) were solubilized on ice with a probe sonicator with 30% amplitude in 5% SDS and 50 mM TEAB pH 8.5. Cycles of 10 seconds on and 10 seconds off were used for 1 minute. Protein concentration was assessed using the Pierce BCA Protein Assay Kit (Thermo Fisher Scientific) with 25 µg of each sample aliquoted in triplicates. 20 µg of whole cell pellets were prepared for mass spectrometry analysis with S-Trap™ mini spin columns (ProtiFi) as per manufacturer’s instructions. With the S-Trap™ micro column in a 1.7 mL tube for flow through, the acidified SDS lysate/MeOH S-Trap™ buffer mixture was added into the micro column. The micro column was then centrifuged at 6500 rpm for 30 s until all SDS lysate/ S-Trap™ buffer had passed through the S-Trap™ column. The captured protein was then washed with 350 µL S-Trap™ buffer with centrifugation and washing repeated three times. The spin column was then transferred to a fresh 1.7 mL tube. Proteins were digested with trypsin (Thermo Fisher Scientific) at 1:10 trypsin to protein ratio and incubated overnight at 37°C. Peptides were eluted with 80 µL each of digestion buffer (50 mM TEAB) and then 0.2% aqueous formic acid was added to the S-Trap™ protein trapping matrix centrifuged at 3600 rpm for 60 s for each elution.

Further peptide purification was performed using SDB-RPS disc stage tips. Peptides were then centrifuged through the column at 1500 g for 3 min. Stage tips were washed with 100 uL of 90% isopropanol (C_3_H_8_O) containing 1% TFA and then washed again with 0.2% TFA in 5% acetonitrile and centrifuged for 4 min following each wash. Peptides were then eluted by adding 100 uL of 60% acetonitrile containing 5% ammonium hydroxide and centrifuged for 4 min. Peptides were dried down using a CentriVap Benchtop Vacuum Concentrator (Labconco). For LC-MS/MS, 2 µL was injected and samples were analyzed on a 170 min gradient at at 250 nL/min and at 40 °C on an Orbitrap Eclipse mass spectrometer (Thermo Fisher Scientific) operating in data-independent acquisition (DIA) mode.

As per the described protocol above, a skeletal muscle library made up of 12 fractionated samples was generated for the search of data-independent acquisition (DIA) of the skeletal muscle samples. For the library, samples were analyzed for 170 minutes on an Orbitrap Eclipse mass spectrometer (Thermo Fisher Scientific) operating in data-dependent acquisition (DDA) mode. For the generation of the library, raw files were imported into Spectronaut (Bruderer et al., 2015)(v.15.2.210819.50606) and the library was generated using the ‘Pulsar’ option with default BGS Factory settings, searching against Uniprot human database (42,386 entries). The resulting library contained 54,900 precursors used for the downstream DIA analysis.

Raw data analysis on the samples was conducted with Spectronaut (Bruderer *et al*., 2015)(v.15.2.210819.50606) against the DDA library generated above. Default BGS Factory search parameters were used with the data filtering setting set to ‘Q-value sparse’ with no imputation during this analysis phase. Oxidation of methionine and N-terminal acetylation were specified as variable modifications, and carbamidomethylation of cysteine was set as a fixed modification. Raw data will be uploaded to the PRIDE repository upon publication.

### Statistics

All individual values are shown together with the group mean. In time-course figures, the data is represented as mean ± 95% confidence interval. Normality was assessed with a Shapiro-Wilk test, and datasets were log-transformed if they failed the normality test. To investigate the effect of exercise (5 time-points) and group (MICE or SIE), and the interaction between these, two-way repeated measures of ANOVA were performed. Interactions were followed by Sidak post-hoc tests to assess the differences between time points (both within and between groups). Main effects of exercise were further analyzed with one-way ANOVA pre-planned contrasts to compare the effects of exercise or training within groups. Due to the extent of data skewness, and inability to normalize mitochondrial morphology variables, these were analyzed with non-parametric Mann-Whitney tests. For RNA-sequencing and proteomics, an adjusted p value (p < 0.05) was used to establish statistical significance. For geneset enrichment analysis the false discovery rate (FDR) q-value was used to set statistical significance (< 0.05). The level of statistical significance was set at p < 0.05. GraphPad Prism 8.3 software was used for all statistical analyses.

## Author Contributions Statement

J.B. and D.J.B. conceptualised the study. J.B., N.A.J., V.O., G.R., A.G., E.F.V., C.U., D.S., M.L. and D.J.B. designed and established the methods. J.B., E.P., N.A.J., J.D.L., N.S., and A.G. carried out the training study and sample collection. J.B., E.P., N.J.C., S.L.C., M.B., N.A.J., V.O., N.S., D.F.T., E.F.V. and C.U. performed the experiments. J.B. and D.J.B. wrote the manuscript. All authors edited and revised the manuscript. All persons designated as authors qualify for authorship, and all those qualifying for authorship are listed. All authors have read and approved the final manuscript.

## Acknowledgments

We wish to thank all the participants for their time and commitment in the present study. The authors would also like to thank the technical team at Victoria University for their assistance throughout the study. We thank the members of the Bishop lab and the Stroud lab for their assistance during the study. We thank the Bio21 Mass Spectrometry and Proteomics Facility (MMSPF) for the provision of instrumentation, training, and technical support.

## REFERENCES

Adhihetty, P.J., Irrcher, I., Joseph, A.M., Ljubicic, V., and Hood, D.A. (2003). Plasticity of skeletal muscle mitochondria in response to contractile activity. Exp Physiol 88, 99–107. 10.1113/eph8802505.

Amar, D., Lindholm, M.E., Norrbom, J., Wheeler, M.T., Rivas, M.A., and Ashley, E.A. (2021). Time trajectories in the transcriptomic response to exercise - a meta-analysis. Nature Communications 12, 3471. 10.1038/s41467-021-23579-x.

Balsa, E., Soustek, M.S., Thomas, A., Cogliati, S., García-Poyatos, C., Martín-García, E., Jedrychowski, M., Gygi, S.P., Enriquez, J.A., and Puigserver, P. (2019). ER and Nutrient Stress Promote Assembly of Respiratory Chain Supercomplexes through the PERK-eIF2&#x3b1; Axis. Molecular Cell 74, 877–890.e876. 10.1016/j.molcel.2019.03.031.

Bao, X.R., Ong, S.-E., Goldberger, O., Peng, J., Sharma, R., Thompson, D.A., Vafai, S.B., Cox, A.G., Marutani, E., Ichinose, F., et al. (2016). Mitochondrial dysfunction remodels one-carbon metabolism in human cells. eLife 5, e10575. 10.7554/eLife.10575.

Bartlett, J.D., Close, G.L., MacLaren, D.P., Gregson, W., Drust, B., and Morton, J.P. (2011). High-intensity interval running is perceived to be more enjoyable than moderate-intensity continuous exercise: implications for exercise adherence. J Sports Sci 29, 547–553. 10.1080/02640414.2010.545427.

Benegiamo, G., Bou Sleiman, M., Wohlwend, M., Rodríguez-López, S., Goeminne, L.J.E., Laurila, P.-P., Klevjer, M., Salonen, M.K., Lahti, J., Jha, P., et al. (2022). COX7A2L genetic variants determine cardiorespiratory fitness in mice and human. Nature Metabolism 4, 1336–1351. 10.1038/s42255-022-00655-0.

Bishop, D.J., Botella, J., Genders, A.J., Lee, M.J., Saner, N.J., Kuang, J., Yan, X., and Granata, C. (2019a). High-Intensity Exercise and Mitochondrial Biogenesis: Current Controversies and Future Research Directions. Physiology (Bethesda, Md.) 34, 56–70. 10.1152/physiol.00038.2018.

Bishop, D.J., Botella, J., and Granata, C. (2019b). CrossTalk opposing view: Exercise training volume is more important than training intensity to promote increases in mitochondrial content. J Physiol 597, 4115–4118. 10.1113/jp277634.

Bishop, D.J., Lee, M.J., and Picard, M. (2024). Exercise as Mitochondrial Medicine: How Does the Exercise Prescription Affect Mitochondrial Adaptations to Training? Annual review of physiology. 10.1146/annurev-physiol-022724-104836.

Blazev, R., Carl, C.S., Ng, Y.-K., Molendijk, J., Voldstedlund, C.T., Zhao, Y., Xiao, D., Kueh, A.J., Miotto, P.M., Haynes, V.R., et al. (2022). Phosphoproteomics of three exercise modalities identifies canonical signaling and C18ORF25 as an AMPK substrate regulating skeletal muscle function. Cell Metabolism. 10.1016/j.cmet.2022.07.003.

Booth, F.W., Roberts, C.K., and Laye, M.J. (2012). Lack of exercise is a major cause of chronic diseases. Comprehensive Physiology 2, 1143.

Botella, J., Motanova, E.S., and Bishop, D.J. (2022). Muscle contraction and mitochondrial biogenesis - A brief historical reappraisal. Acta Physiol (Oxf) 235, e13813. 10.1111/apha.13813.

Botella, J., Schytz, C.T., Pehrson, T.F., Hokken, R., Laugesen, S., Aagaard, P., Suetta, C., Christensen, B., Ørtenblad, N., and Nielsen, J. (2023a). Increased mitochondrial surface area and cristae density in the skeletal muscle of strength athletes. J Physiol 601, 2899–2915. 10.1113/jp284394.

Botella, J., Shaw, C.S., and Bishop, D.J. (2023b). Autophagy and Exercise: Current Insights and Future Research Directions. Int J Sports Med. 10.1055/a-2153-9258.

Brischigliaro, M., Cabrera-Orefice, A., Arnold, S., Viscomi, C., Zeviani, M., and Fernández-Vizarra, E. (2023). Structural rather than catalytic role for mitochondrial respiratory chain supercomplexes. eLife Sciences Publications, Ltd.

Brischigliaro, M., Frigo, E., Fernandez-Vizarra, E., Bernardi, P., and Viscomi, C. (2022). Measurement of mitochondrial respiratory chain enzymatic activities in Drosophila melanogaster samples. STAR Protoc 3, 101322. 10.1016/j.xpro.2022.101322.

Broskey, N.T., Daraspe, J., Humbel, B.M., and Amati, F. (2013). Skeletal muscle mitochondrial and lipid droplet content assessed with standardized grid sizes for stereology. Journal of Applied Physiology 115, 765–770. 10.1152/japplphysiol.00063.2013.

Bruderer, R., Bernhardt, O.M., Gandhi, T., Miladinović, S.M., Cheng, L.Y., Messner, S., Ehrenberger, T., Zanotelli, V., Butscheid, Y., Escher, C., et al. (2015). Extending the limits of quantitative proteome profiling with data-independent acquisition and application to acetaminophen-treated three-dimensional liver microtissues. Mol Cell Proteomics 14, 1400–1410. 10.1074/mcp.M114.044305.

Burgomaster, K.A., Howarth, K.R., Phillips, S.M., Rakobowchuk, M., Macdonald, M.J., McGee, S.L., and Gibala, M.J. (2008). Similar metabolic adaptations during exercise after low volume sprint interval and traditional endurance training in humans. J Physiol 586, 151–160. 10.1113/jphysiol.2007.142109.

Callegari, S., Richter, F., Chojnacka, K., Jans, D.C., Lorenzi, I., Pacheu-Grau, D., Jakobs, S., Lenz, C., Urlaub, H., Dudek, J., et al. (2016). TIM29 is a subunit of the human carrier translocase required for protein transport. FEBS Lett 590, 4147–4158. 10.1002/1873-3468.12450.

Cardinale, D.A., Gejl, K.D., Petersen, K.G., Nielsen, J., Ørtenblad, N., and Larsen, F.J. (2021). Short-term intensified training temporarily impairs mitochondrial respiratory capacity in elite endurance athletes. Journal of Applied Physiology 131, 388–400. 10.1152/japplphysiol.00829.2020.

Cochran, A.J.R., Percival, M.E., Tricarico, S., Little, J.P., Cermak, N., Gillen, J.B., Tarnopolsky, M.A., and Gibala, M.J. (2014). Intermittent and continuous high-intensity exercise training induce similar acute but different chronic muscle adaptations. Experimental Physiology 99, 782–791. 10.1113/expphysiol.2013.077453.

Cordeiro, A.V., Brícola, R.S., Braga, R.R., Lenhare, L., Silva, V.R.R., Anaruma, C.P., Katashima, C.K., Crisol, B.M., Simabuco, F.M., Silva, A.S.R., et al. (2020). Aerobic Exercise Training Induces the Mitonuclear Imbalance and UPRmt in the Skeletal Muscle of Aged Mice. J Gerontol A Biol Sci Med Sci 75, 2258–2261. 10.1093/gerona/glaa059.

Cordeiro, A.V., Peruca, G.F., Braga, R.R., Brícola, R.S., Lenhare, L., Silva, V.R.R., Anaruma, C.P., Katashima, C.K., Crisol, B.M., Barbosa, L.T., et al. (2021). High-intensity exercise training induces mitonuclear imbalance and activates the mitochondrial unfolded protein response in the skeletal muscle of aged mice. Geroscience 43, 1513–1518. 10.1007/s11357-020-00246-5.

Escobar-Alvarez, S., Gardner, J., Sheth, A., Manfredi, G., Yang, G., Ouerfelli, O., Heaney, M.L., and Scheinberg, D.A. (2010). Inhibition of Human Peptide Deformylase Disrupts Mitochondrial Function. Molecular and Cellular Biology 30, 5099–5109. 10.1128/MCB.00469-10.

Fernández-Vizarra, E., López-Calcerrada, S., Sierra-Magro, A., Pérez-Pérez, R., Formosa, L.E., Hock, D.H., Illescas, M., Peñas, A., Brischigliaro, M., Ding, S., et al. (2022). Two independent respiratory chains adapt OXPHOS performance to glycolytic switch. Cell Metab 34, 1792–1808.e1796. 10.1016/j.cmet.2022.09.005.

Fernández-Vizarra, E., and Ugalde, C. (2022). Cooperative assembly of the mitochondrial respiratory chain. Trends in Biochemical Sciences 47, 999–1008. 10.1016/j.tibs.2022.07.005.

Fessler, E., Eckl, E.-M., Schmitt, S., Mancilla, I.A., Meyer-Bender, M.F., Hanf, M., Philippou-Massier, J., Krebs, S., Zischka, H., and Jae, L.T. (2020). A pathway coordinated by DELE1 relays mitochondrial stress to the cytosol. Nature 579, 433–437. 10.1038/s41586-020-2076-4.

Flockhart, M., Nilsson, L.C., Tais, S., Ekblom, B., Apró, W., and Larsen, F.J. (2021). Excessive exercise training causes mitochondrial functional impairment and decreases glucose tolerance in healthy volunteers. Cell Metabolism 33, 957–970.e956. 10.1016/j.cmet.2021.02.017.

Forsström, S., Jackson, C.B., Carroll, C.J., Kuronen, M., Pirinen, E., Pradhan, S., Marmyleva, A., Auranen, M., Kleine, I.M., Khan, N.A., et al. (2019). Fibroblast Growth Factor 21 Drives Dynamics of Local and Systemic Stress Responses in Mitochondrial Myopathy with mtDNA Deletions. Cell Metab 30, 1040–1054.e1047. 10.1016/j.cmet.2019.08.019.

Garber, C.E., Blissmer, B., Deschenes, M.R., Franklin, B.A., Lamonte, M.J., Lee, I.-M., Nieman, D.C., and Swain, D.P. (2011). Quantity and Quality of Exercise for Developing and Maintaining Cardiorespiratory, Musculoskeletal, and Neuromotor Fitness in Apparently Healthy Adults: Guidance for Prescribing Exercise. Medicine & Science in Sports & Exercise 43, 1334–1359. 10.1249/MSS.0b013e318213fefb.

Gibala, M.J., and Hawley, J.A. (2017). Sprinting Toward Fitness. Cell Metab 25, 988–990. 10.1016/j.cmet.2017.04.030.

Gillen, J.B., Martin, B.J., MacInnis, M.J., Skelly, L.E., Tarnopolsky, M.A., and Gibala, M.J. (2016). Twelve Weeks of Sprint Interval Training Improves Indices of Cardiometabolic Health Similar to Traditional Endurance Training despite a Five-Fold Lower Exercise Volume and Time Commitment. PloS one 11, e0154075. 10.1371/journal.pone.0154075.

Glancy, B., Hartnell, L.M., Combs, C.A., Femnou, A., Sun, J., Murphy, E., Subramaniam, S., and Balaban, R.S. (2017). Power Grid Protection of the Muscle Mitochondrial Reticulum. Cell Rep 19, 487–496. 10.1016/j.celrep.2017.03.063.

Godin, G., Desharnais, R., Valois, P., Lepage, L., Jobin, J., and Bradet, R. (1994). Differences in Perceived Barriers to Exercise between High and Low Intenders: Observations among Different Populations. American Journal of Health Promotion 8, 279–285. 10.4278/0890-1171-8.4.279.

Gollnick, P., and King, D. (1969). Effect of exercise and training on mitochondria of rat skeletal muscle. American Journal of Physiology-Legacy Content 216, 1502–1509. 10.1152/ajplegacy.1969.216.6.1502.

Gollnick, P.D., and King, D.W. (1971). Advances in Experimental Medicine and Biology. pp. 69–85 p.

Granata, C., Caruana, N.J., Botella, J., Jamnick, N.A., Huynh, K., Kuang, J., Janssen, H.A., Reljic, B., Mellett, N.A., Laskowski, A., et al. (2021). High-intensity training induces non-stoichiometric changes in the mitochondrial proteome of human skeletal muscle without reorganisation of respiratory chain content. Nature Communications 12, 7056. 10.1038/s41467-021-27153-3.

Granata, C., Jamnick, N.A., and Bishop, D.J. (2018). Training-Induced Changes in Mitochondrial Content and Respiratory Function in Human Skeletal Muscle. Sports medicine (Auckland, N.Z.) 48, 1809–1828. 10.1007/s40279-018-0936-y.

Granata, C., Oliveira, R.S., Little, J.P., Renner, K., and Bishop, D.J. (2016a). Training intensity modulates changes in PGC-1alpha and p53 protein content and mitochondrial respiration, but not markers of mitochondrial content in human skeletal muscle. FASEB journal : official publication of the Federation of American Societies for Experimental Biology 30, 959–970. 10.1096/fj.15-276907.

Granata, C., Oliveira, R.S.F., Little, J.P., Renner, K., and Bishop, D.J. (2016b). Mitochondrial adaptations to high-volume exercise training are rapidly reversed after a reduction in training volume in human skeletal muscle. The FASEB Journal 30, 3413–3423. 10.1096/fj.201500100R.

Granata, C., Oliveira, R.S.F., Little, J.P., Renner, K., and Bishop, D.J. (2017). Sprint-interval but not continuous exercise increases PGC-1α protein content and p53 phosphorylation in nuclear fractions of human skeletal muscle. Scientific Reports 7, 44227. 10.1038/srep44227.

Greggio, C., Jha, P., Kulkarni, S.S., Lagarrigue, S., Broskey, N.T., Boutant, M., Wang, X., Conde Alonso, S., Ofori, E., Auwerx, J., et al. (2017). Enhanced Respiratory Chain Supercomplex Formation in Response to Exercise in Human Skeletal Muscle. Cell Metab 25, 301–311. 10.1016/j.cmet.2016.11.004.

Grevendonk, L., Connell, N.J., McCrum, C., Fealy, C.E., Bilet, L., Bruls, Y.M.H., Mevenkamp, J., Schrauwen-Hinderling, V.B., Jörgensen, J.A., Moonen-Kornips, E., et al. (2021). Impact of aging and exercise on skeletal muscle mitochondrial capacity, energy metabolism, and physical function. Nature Communications 12, 4773. 10.1038/s41467-021-24956-2.

Guo, X., Aviles, G., Liu, Y., Tian, R., Unger, B.A., Lin, Y.-H.T., Wiita, A.P., Xu, K., Correia, M.A., and Kampmann, M. (2020). Mitochondrial stress is relayed to the cytosol by an OMA1–DELE1–HRI pathway. Nature 579, 427–432. 10.1038/s41586-020-2078-2.

Haskell, W.L., Lee, I.M., Pate, R.R., Powell, K.E., Blair, S.N., Franklin, B.A., Macera, C.A., Heath, G.W., Thompson, P.D., and Bauman, A. (2007). Physical activity and public health: updated recommendation for adults from the American College of Sports Medicine and the American Heart Association. Circulation 116, 1081–1093. 10.1161/circulationaha.107.185649.

Helge, J., Damsgaard, R., Overgaard, K., Andersen, J., Donsmark, M., Dyrskog, S., Hermansen, K., Saltin, B., and Daugaard, J. (2008). Low-intensity training dissociates metabolic from aerobic fitness. Scand. J. Med. Sci. Sports 18, 86–94.

Heo, J.M., Ordureau, A., Paulo, J.A., Rinehart, J., and Harper, J.W. (2015). The PINK1-PARKIN Mitochondrial Ubiquitylation Pathway Drives a Program of OPTN/NDP52 Recruitment and TBK1 Activation to Promote Mitophagy. Mol Cell 60, 7–20. 10.1016/j.molcel.2015.08.016.

Holloszy, J.O. (1967). Biochemical adaptations in muscle. Effects of exercise on mitochondrial oxygen uptake and respiratory enzyme activity in skeletal muscle. The Journal of biological chemistry 242, 2278–2282.

Hood, D.A., Memme, J.M., Oliveira, A.N., and Triolo, M. (2019). Maintenance of Skeletal Muscle Mitochondria in Health, Exercise, and Aging. Annual review of physiology 81, 19–41. 10.1146/annurev-physiol-020518-114310.

Hood, D.A., Uguccioni, G., Vainshtein, A., and D’Souza, D. (2011). Mechanisms of exercise-induced mitochondrial biogenesis in skeletal muscle: implications for health and disease. Comprehensive Physiology 1, 1119–1134. 10.1002/cphy.c100074.

Hoppeler, H., Lüthi, P., Claassen, H., Weibel, E.R., and Howald, H. (1973). The ultrastructure of the normal human skeletal muscle. A morphometric analysis on untrained men, women and well-trained orienteers. Pflugers Arch 344, 217–232. 10.1007/bf00588462.

Hostrup, M., Lemminger, A.K., Stocks, B., Gonzalez-Franquesa, A., Larsen, J.K., Quesada, J.P., Thomassen, M., Weinert, B.T., Bangsbo, J., and Deshmukh, A.S. (2022). High-intensity interval training remodels the proteome and acetylome of human skeletal muscle. eLife 11, e69802. 10.7554/eLife.69802.

Houzelle, A., Jörgensen, J.A., Schaart, G., Daemen, S., van Polanen, N., Fealy, C.E., Hesselink, M.K.C., Schrauwen, P., and Hoeks, J. (2021). Human skeletal muscle mitochondrial dynamics in relation to oxidative capacity and insulin sensitivity. Diabetologia 64, 424–436. 10.1007/s00125-020-05335-w.

Huertas, J.R., Ruiz-Ojeda, F.J., Plaza-Díaz, J., Nordsborg, N.B., Martín-Albo, J., Rueda-Robles, A., and Casuso, R.A. (2019). Human muscular mitochondrial fusion in athletes during exercise. The FASEB Journal 33, 12087–12098. 10.1096/fj.201900365RR.

Jacobs, R.A., and Lundby, C. (2013). Mitochondria express enhanced quality as well as quantity in association with aerobic fitness across recreationally active individuals up to elite athletes. J Appl Physiol (1985) 114, 344–350. 10.1152/japplphysiol.01081.2012.

Jamnick, N.A., Botella, J., Pyne, D.B., and Bishop, D.J. (2018). Manipulating graded exercise test variables affects the validity of the lactate threshold and V. O2peak. PLoS One 13, e0199794.

Jamnick, N.A., Pettitt, R.W., Granata, C., Pyne, D.B., and Bishop, D.J. (2020). An Examination and Critique of Current Methods to Determine Exercise Intensity. Sports Medicine 50, 1729–1756. 10.1007/s40279-020-01322-8.

Kaspi, A., and Ziemann, M. (2020). mitch: multi-contrast pathway enrichment for multi-omics and single-cell profiling data. BMC Genomics 21, 447. 10.1186/s12864-020-06856-9.

Khan, N.A., Nikkanen, J., Yatsuga, S., Jackson, C., Wang, L., Pradhan, S., Kivelä, R., Pessia, A., Velagapudi, V., and Suomalainen, A. (2017). mTORC1 Regulates Mitochondrial Integrated Stress Response and Mitochondrial Myopathy Progression. Cell Metabolism 26, 419–428.e415. 10.1016/j.cmet.2017.07.007.

Kleele, T., Rey, T., Winter, J., Zaganelli, S., Mahecic, D., Perreten Lambert, H., Ruberto, F.P., Nemir, M., Wai, T., Pedrazzini, T., and Manley, S. (2021). Distinct fission signatures predict mitochondrial degradation or biogenesis. Nature 593, 435–439. 10.1038/s41586-021-03510-6.

Korotkevich, G., Sukhov, V., Budin, N., Shpak, B., Artyomov, M.N., and Sergushichev, A. (2021). Fast gene set enrichment analysis. bioRxiv, 060012. 10.1101/060012.

Kuang, J., McGinley, C., Lee, M.J., Saner, N.J., Garnham, A., and Bishop, D.J. (2022). Interpretation of exercise-induced changes in human skeletal muscle mRNA expression depends on the timing of the post-exercise biopsies. PeerJ 10, e12856.

Kuang, J., Yan, X., Genders, A.J., Granata, C., and Bishop, D.J. (2018). An overview of technical considerations when using quantitative real-time PCR analysis of gene expression in human exercise research. PloS one 13, e0196438. 10.1371/journal.pone.0196438.

Labbé, K., LeBon, L., King, B., Vu, N., Stoops, E.H., Ly, N., Lefebvre, A.E.Y.T., Seitzer, P., Krishnan, S., Heo, J.-M., et al. (2024). Specific activation of the integrated stress response uncovers regulation of central carbon metabolism and lipid droplet biogenesis. Nature Communications 15, 8301. 10.1038/s41467-024-52538-5.

Laker, R.C., Drake, J.C., Wilson, R.J., Lira, V.A., Lewellen, B.M., Ryall, K.A., Fisher, C.C., Zhang, M., Saucerman, J.J., Goodyear, L.J., et al. (2017). Ampk phosphorylation of Ulk1 is required for targeting of mitochondria to lysosomes in exercise-induced mitophagy. Nat Commun 8, 548. 10.1038/s41467-017-00520-9.

Lavorato, M., Loro, E., Debattisti, V., Khurana, T.S., and Franzini-Armstrong, C. (2018). Elongated mitochondrial constrictions and fission in muscle fatigue. Journal of Cell Science 131, jcs221028. 10.1242/jcs.221028.

Lazarou, M., Sliter, D.A., Kane, L.A., Sarraf, S.A., Wang, C., Burman, J.L., Sideris, D.P., Fogel, A.I., and Youle, R.J. (2015). The ubiquitin kinase PINK1 recruits autophagy receptors to induce mitophagy. Nature 524, 309–314. 10.1038/nature14893.

Lee, I.-M., Hsieh, C.-c., and Paffenbarger, R.S., Jr (1995). Exercise Intensity and Longevity in Men: The Harvard Alumni Health Study. JAMA 273, 1179–1184. 10.1001/jama.1995.03520390039030.

Lee, I.M., Shiroma, E.J., Lobelo, F., Puska, P., Blair, S.N., and Katzmarzyk, P.T. (2012). Effect of physical inactivity on major non-communicable diseases worldwide: an analysis of burden of disease and life expectancy. Lancet (London, England) 380, 219–229. 10.1016/s0140-6736(12)61031-9.

Linares, J.F., Duran, A., Reina-Campos, M., Aza-Blanc, P., Campos, A., Moscat, J., and Diaz-Meco, M.T. (2015). Amino Acid Activation of mTORC1 by a PB1-Domain-Driven Kinase Complex Cascade. Cell Rep 12, 1339–1352. 10.1016/j.celrep.2015.07.045.

Lovrić, A., Rassolie, A., Alam, S., Mandić, M., Saini, A., Altun, M., Fernandez-Gonzalo, R., Gustafsson, T., and Rullman, E. (2022). Single-cell sequencing deconvolutes cellular responses to exercise in human skeletal muscle. Communications Biology 5, 1121. 10.1038/s42003-022-04088-z.

MacInnis, M.J., Skelly, L.E., and Gibala, M.J. (2019). CrossTalk proposal: Exercise training intensity is more important than volume to promote increases in human skeletal muscle mitochondrial content. J Physiol 597, 4111–4113. 10.1113/jp277633.

Meinild Lundby, A.K., Jacobs, R.A., Gehrig, S., de Leur, J., Hauser, M., Bonne, T.C., Flück, D., Dandanell, S., Kirk, N., Kaech, A., et al. (2018). Exercise training increases skeletal muscle mitochondrial volume density by enlargement of existing mitochondria and not de novo biogenesis. Acta Physiol (Oxf) 222. 10.1111/apha.12905.

Melber, A., and Haynes, C.M. (2018). UPR(mt) regulation and output: a stress response mediated by mitochondrial-nuclear communication. Cell Res 28, 281–295. 10.1038/cr.2018.16.

Merry, T.L., and Ristow, M. (2016). Mitohormesis in exercise training. Free Radic Biol Med 98, 123–130. 10.1016/j.freeradbiomed.2015.11.032.

Migliavacca, E., Tay, S.K.H., Patel, H.P., Sonntag, T., Civiletto, G., McFarlane, C., Forrester, T., Barton, S.J., Leow, M.K., Antoun, E., et al. (2019). Mitochondrial oxidative capacity and NAD+ biosynthesis are reduced in human sarcopenia across ethnicities. Nature Communications 10, 5808. 10.1038/s41467-019-13694-1.

Milenkovic, D., Misic, J., Hevler, J.F., Molinié, T., Chung, I., Atanassov, I., Li, X., Filograna, R., Mesaros, A., Mourier, A., et al. (2023). Preserved respiratory chain capacity and physiology in mice with profoundly reduced levels of mitochondrial respirasomes. Cell Metab 35, 1799–1813.e1797. 10.1016/j.cmet.2023.07.015.

Neufer, P.D., Bamman, M.M., Muoio, D.M., Bouchard, C., Cooper, D.M., Goodpaster, B.H., Booth, F.W., Kohrt, W.M., Gerszten, R.E., Mattson, M.P., et al. (2015). Understanding the Cellular and Molecular Mechanisms of Physical Activity-Induced Health Benefits. Cell Metab 22, 4–11. 10.1016/j.cmet.2015.05.011.

Nielsen, J., Gejl, K.D., Hey-Mogensen, M., Holmberg, H.C., Suetta, C., Krustrup, P., Elemans, C.P.H., and Ørtenblad, N. (2017). Plasticity in mitochondrial cristae density allows metabolic capacity modulation in human skeletal muscle. J Physiol 595, 2839–2847. 10.1113/jp273040.

Nimmo, M.A., and Snow, D.H. (1982). Time course of ultrastructural changes in skeletal muscle after two types of exercise. J Appl Physiol Respir Environ Exerc Physiol 52, 910–913. 10.1152/jappl.1982.52.4.910.

Pakos-Zebrucka, K., Koryga, I., Mnich, K., Ljujic, M., Samali, A., and Gorman, A.M. (2016). The integrated stress response. EMBO reports 17, 1374–1395. 10.15252/embr.201642195.

Parker, L., Trewin, A., Levinger, I., Shaw, C.S., and Stepto, N.K. (2017). The effect of exercise-intensity on skeletal muscle stress kinase and insulin protein signaling. PloS one 12, e0171613–e0171613. 10.1371/journal.pone.0171613.

Perry, C.G.R., Lally, J., Holloway, G.P., Heigenhauser, G.J.F., Bonen, A., and Spriet, L.L. (2010). Repeated transient mRNA bursts precede increases in transcriptional and mitochondrial proteins during training in human skeletal muscle. J Physiol 588, 4795–4810. 10.1113/jphysiol.2010.199448.

Pfaffl, M.W., Tichopad, A., Prgomet, C., and Neuvians, T.P. (2004). Determination of stable housekeeping genes, differentially regulated target genes and sample integrity: BestKeeper – Excel-based tool using pair-wise correlations. Biotechnology Letters 26, 509–515. 10.1023/B:BILE.0000019559.84305.47.

Picard, M., Gentil, B.J., McManus, M.J., White, K., St Louis, K., Gartside, S.E., Wallace, D.C., and Turnbull, D.M. (2013). Acute exercise remodels mitochondrial membrane interactions in mouse skeletal muscle. J Appl Physiol (1985) 115, 1562–1571. 10.1152/japplphysiol.00819.2013.

Pillon, N.J., Gabriel, B.M., Dollet, L., Smith, J.A.B., Sardón Puig, L., Botella, J., Bishop, D.J., Krook, A., and Zierath, J.R. (2020). Transcriptomic profiling of skeletal muscle adaptations to exercise and inactivity. Nature Communications 11, 470. 10.1038/s41467-019-13869-w.

Place, N., Ivarsson, N., Venckunas, T., Neyroud, D., Brazaitis, M., Cheng, A.J., Ochala, J., Kamandulis, S., Girard, S., Volungevicius, G., et al. (2015). Ryanodine receptor fragmentation and sarcoplasmic reticulum Ca2+ leak after one session of high-intensity interval exercise. Proceedings of the National Academy of Sciences of the United States of America 112, 15492–15497. 10.1073/pnas.1507176112.

Quiros, P.M., Prado, M.A., Zamboni, N., D’Amico, D., Williams, R.W., Finley, D., Gygi, S.P., and Auwerx, J. (2017). Multi-omics analysis identifies ATF4 as a key regulator of the mitochondrial stress response in mammals. The Journal of cell biology 216, 2027–2045. 10.1083/jcb.201702058.

Rath, S., Sharma, R., Gupta, R., Ast, T., Chan, C., Durham, T.J., Goodman, R.P., Grabarek, Z., Haas, M.E., Hung, W.H.W., et al. (2021). MitoCarta3.0: an updated mitochondrial proteome now with sub-organelle localization and pathway annotations. Nucleic Acids Res 49, D1541–d1547. 10.1093/nar/gkaa1011.

Reisman, E.G., Botella, J., Huang, C., Schittenhelm, R.B., Stroud, D.A., Granata, C., Chandrasiri, O.S., Ramm, G., Oorschot, V., Caruana, N.J., and Bishop, D.J. (2024). Fibre-specific mitochondrial protein abundance is linked to resting and post-training mitochondrial content in the muscle of men. Nature Communications 15, 7677. 10.1038/s41467-024-50632-2.

Richter, B., Sliter, D.A., Herhaus, L., Stolz, A., Wang, C., Beli, P., Zaffagnini, G., Wild, P., Martens, S., Wagner, S.A., et al. (2016). Phosphorylation of OPTN by TBK1 enhances its binding to Ub chains and promotes selective autophagy of damaged mitochondria. Proceedings of the National Academy of Sciences 113, 4039–4044. doi:10.1073/pnas.1523926113.

Robinson, M.M., Dasari, S., Konopka, A.R., Johnson, M.L., Manjunatha, S., Esponda, R.R., Carter, R.E., Lanza, I.R., and Nair, K.S. (2017). Enhanced Protein Translation Underlies Improved Metabolic and Physical Adaptations to Different Exercise Training Modes in Young and Old Humans. Cell Metab 25, 581–592. 10.1016/j.cmet.2017.02.009.

Rowe, G.C., Patten, I.S., Zsengeller, Z.K., El-Khoury, R., Okutsu, M., Bampoh, S., Koulisis, N., Farrell, C., Hirshman, M.F., Yan, Z., et al. (2013). Disconnecting mitochondrial content from respiratory chain capacity in PGC-1-deficient skeletal muscle. Cell Rep 3, 1449–1456. 10.1016/j.celrep.2013.04.023.

Sahlin, K., Tonkonogi, M., and Söderlund, K. (1998). Energy supply and muscle fatigue in humans. Acta Physiol Scand 162, 261–266. 10.1046/j.1365-201X.1998.0298f.x.

Shpilka, T., and Haynes, C.M. (2018). The mitochondrial UPR: mechanisms, physiological functions and implications in ageing. Nature Reviews Molecular Cell Biology 19, 109–120. 10.1038/nrm.2017.110.

Spinelli, J.B., and Haigis, M.C. (2018). The multifaceted contributions of mitochondria to cellular metabolism. Nat Cell Biol 20, 745–754. 10.1038/s41556-018-0124-1.

Sprenger, H.G., and Langer, T. (2019). The Good and the Bad of Mitochondrial Breakups. Trends Cell Biol 29, 888–900. 10.1016/j.tcb.2019.08.003.

Sutandy, F.X.R., Gößner, I., Tascher, G., and Münch, C. (2023). A cytosolic surveillance mechanism activates the mitochondrial UPR. Nature 618, 849–854. 10.1038/s41586-023-06142-0.

Timón-Gómez, A., Pérez-Pérez, R., Nyvltova, E., Ugalde, C., Fontanesi, F., and Barrientos, A. (2020). Protocol for the Analysis of Yeast and Human Mitochondrial Respiratory Chain Complexes and Supercomplexes by Blue Native Electrophoresis. STAR Protocols 1, 100089. 10.1016/j.xpro.2020.100089.

van Loon, L.J.C., Greenhaff, P.L., Constantin-Teodosiu, D., Saris, W.H.M., and Wagenmakers, A.J.M. (2001). The effects of increasing exercise intensity on muscle fuel utilisation in humans. J Physiol 536, 295–304. 10.1111/j.1469-7793.2001.00295.x.

Vercellino, I., and Sazanov, L.A. (2021). Structure and assembly of the mammalian mitochondrial supercomplex CIII(2)CIV. Nature 598, 364–367. 10.1038/s41586-021-03927-z.

Vercellino, I., and Sazanov, L.A. (2024). SCAF1 drives the compositional diversity of mammalian respirasomes. Nature Structural & Molecular Biology 31, 1061–1071. 10.1038/s41594-024-01255-0.

Wai, T., and Langer, T. (2016). Mitochondrial Dynamics and Metabolic Regulation. Trends in Endocrinology & Metabolism 27, 105–117. 10.1016/j.tem.2015.12.001.

Wang, C., Taki, M., Sato, Y., Tamura, Y., Yaginuma, H., Okada, Y., and Yamaguchi, S. (2019). A photostable fluorescent marker for the superresolution live imaging of the dynamic structure of the mitochondrial cristae. Proceedings of the National Academy of Sciences 116, 15817–15822. doi:10.1073/pnas.1905924116.

Wild, P., Farhan, H., McEwan, D.G., Wagner, S., Rogov, V.V., Brady, N.R., Richter, B., Korac, J., Waidmann, O., Choudhary, C., et al. (2011). Phosphorylation of the autophagy receptor optineurin restricts Salmonella growth. Science 333, 228–233. 10.1126/science.1205405.

Wolf, D.M., Segawa, M., Kondadi, A.K., Anand, R., Bailey, S.T., Reichert, A.S., van der Bliek, A.M., Shackelford, D.B., Liesa, M., and Shirihai, O.S. (2019). Individual cristae within the same mitochondrion display different membrane potentials and are functionally independent. EMBO J 38, e101056. 10.15252/embj.2018101056.

Wong, Y.C., and Holzbaur, E.L. (2014). Optineurin is an autophagy receptor for damaged mitochondria in parkin-mediated mitophagy that is disrupted by an ALS-linked mutation. Proceedings of the National Academy of Sciences of the United States of America 111, E4439–4448. 10.1073/pnas.1405752111.

Wrobel, L., Topf, U., Bragoszewski, P., Wiese, S., Sztolsztener, M.E., Oeljeklaus, S., Varabyova, A., Lirski, M., Chroscicki, P., Mroczek, S., et al. (2015). Mistargeted mitochondrial proteins activate a proteostatic response in the cytosol. Nature 524, 485–488. 10.1038/nature14951.

Wu, J., Ruas, J.L., Estall, J.L., Rasbach, K.A., Choi, J.H., Ye, L., Boström, P., Tyra, H.M., Crawford, R.W., Campbell, K.P., et al. (2011). The unfolded protein response mediates adaptation to exercise in skeletal muscle through a PGC-1α/ATF6α complex. Cell metabolism 13, 160–169. 10.1016/j.cmet.2011.01.003.

Yun, J., and Finkel, T. (2014). Mitohormesis. Cell Metabolism 19, 757–766. 10.1016/j.cmet.2014.01.011.

Zanou, N., Dridi, H., Reiken, S., Imamura de Lima, T., Donnelly, C., De Marchi, U., Ferrini, M., Vidal, J., Sittenfeld, L., , J.N., et al. (2021). Acute RyR1 Ca2+ leak enhances NADH-linked mitochondrial respiratory capacity. Nature Communications 12, 7219. 10.1038/s41467-021-27422-1.

Zhao, Q., Wang, J., Levichkin, I.V., Stasinopoulos, S., Ryan, M.T., and Hoogenraad, N.J. (2002). A mitochondrial specific stress response in mammalian cells. EMBO J 21, 4411–4419. 10.1093/emboj/cdf445.

Zierath, Juleen R., and Wallberg-Henriksson, H. (2015). Looking Ahead Perspective: Where Will the Future of Exercise Biology Take Us? Cell Metabolism 22, 25–30. 10.1016/j.cmet.2015.06.015.

