## Supplemental figures for "Sprint interval exercise disrupts mitochondrial ultrastructure driving a unique mitochondrial stress response and remodelling in humans"

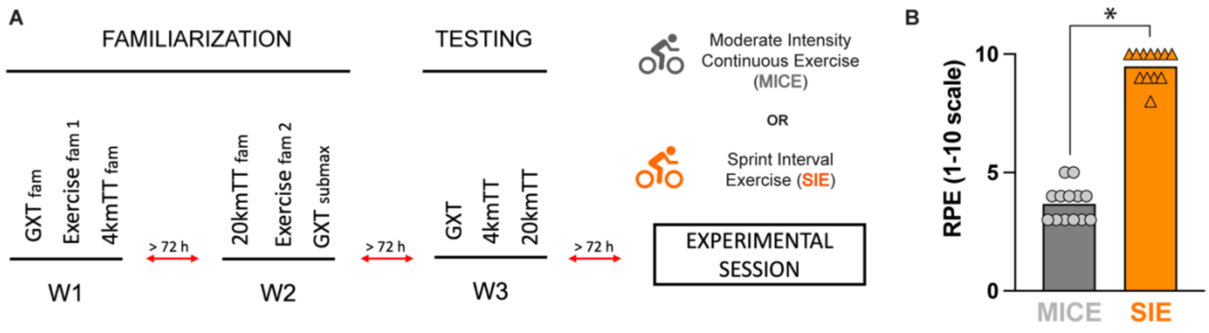


Supplemental Figure 1. A) Schematic of the 3-week familiarization period that participants underwent prior to the experimental session. B) Rating of perceived exertion (RPE) during the experimental session following moderate-intensity continuous exercise (MICE; n = 13) or sprint interval exercise (SIE; n = 12). GXT = graded exercise test; 4kmTT = 4 km time-trial; 20kmTT = 20 km time-trial; fam = familiarization; submax = submaximal. * denotes a main effect at $p < 0.05$. For B) differences were assessed with one-way ANOVA.

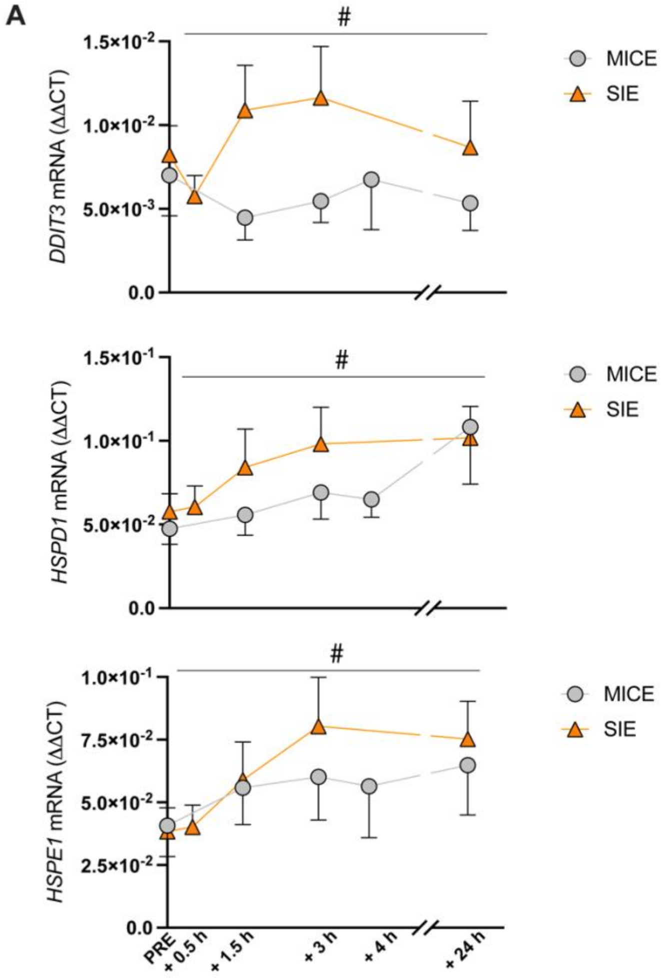


B

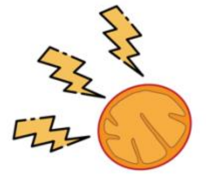


| | NES | FDR |
|---|---|---|
| MICE PRE vs MICE 3 h | 1.06 | 0.33 |
| MICE PRE vs MICE 24 h | 0.90 | 0.62 |
| SIE PRE vs SIE 3 h | **1.50** | **< 0.01** |
| SIE PRE vs SIE 24 h | 0.75 | 0.85 |
| MICE 3 h VS SIE 3 h | 1.27 | 0.10 |

Supplemental Figure 2. A) Changes in the mRNA expression of the mitochondrial unfolded protein response ($UPR^{mt}$) markers with MICE (n = 13) and SIE (n = 14) measures by qPCR. B) Table showing the normalized enrichment score (NES) and false-discovery rate (FDR) of the 'mitostress' geneset in the RNA-sequencing data from both moderate-intensity continuous exercise (MICE) and sprint interval exercise (SIE) across the different timepoints. * denotes main effect at $p < 0.05$; # denotes time x group interaction at $p < 0.05$. For A) data are expressed as mean ± 95% confidence interval. For A) differences were assessed with two-way ANOVA.

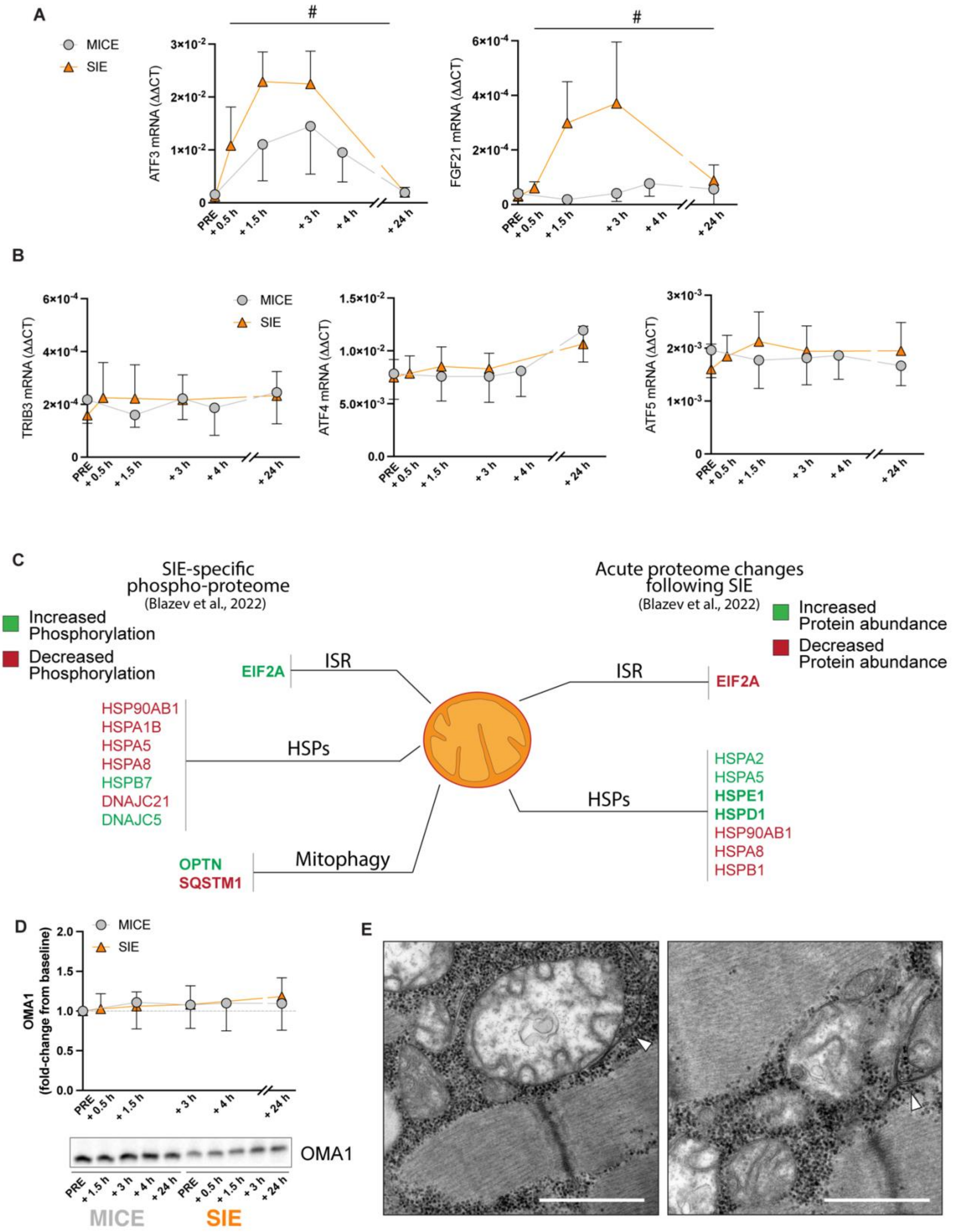


Supplemental Figure 3. A) Changes in mRNA expression of the divergently regulated integrated stress response (ISR) genes ATF3 and FGF21 following moderate-intensity continuous exercise (MICE; n =

13) and sprint interval exercise (SIE; n = 14) measured with qPCR. B) Unchanged mRNA expression of certain genes associated with the ISR following MICE (n = 13) and SIE (n = 14). C) List of proteins that were uniquely post-translationally (phosphorylation status) or quantitatively (protein levels) regulated by SIE in comparison to MICE or resistance exercise (from (Blazev *et al.*, 2022)). In green are proteins phosphorylated and/or increased protein abundance, while red reflects dephosphorylated and/or decreased protein abundance. D) Protein content of OMA1 following MICE (n = 10) and SIE (n = 13). E) Micrographs obtained following SIE (+0h) showing autophagosome-like membranes positioned on the outer membrane of damaged mitochondria. Scale bar = 0.5 µm. # denotes time x group interaction at $p < 0.05$. For A), B), and D) data are expressed as mean ± 95% confidence interval. For A) differences were assessed with two-way ANOVA.

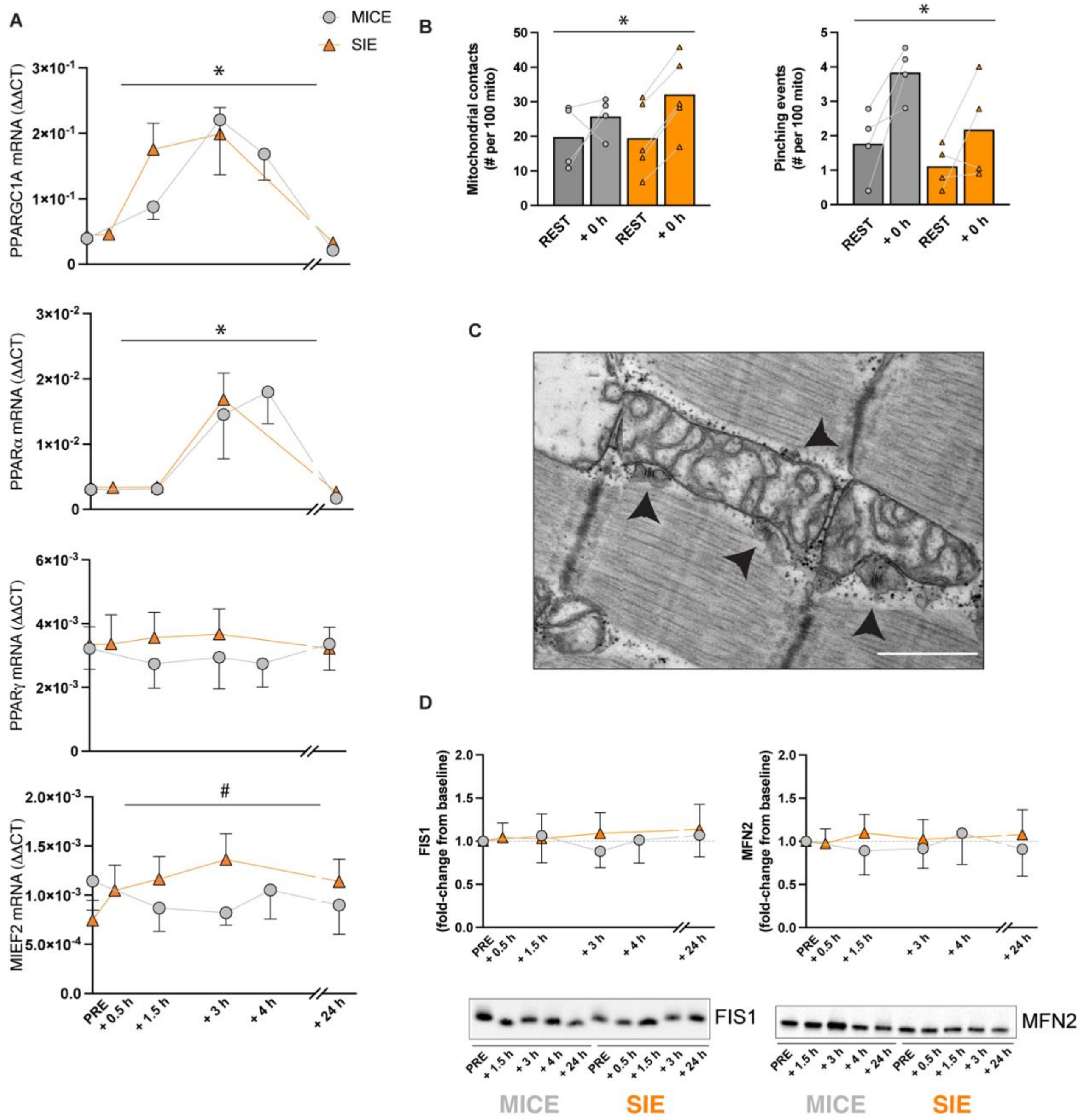


Supplemental Figure 4. A) Changes in the mRNA expression of the selected markers of mitochondrial biogenesis and mitochondrial fission following moderate-intensity continuous exercise (MICE; n = 13) and sprint interval exercise (SIE; n = 14). B) Quantification of mitochondrial contacts (as a pre-fusion event; MICE, n = 4; SIE, n = 5) and mitochondrial pinching (as a pro-fission event; MICE, n = 4; SIE, n = 4) following exercise. C) Micrograph depicting pinching events on the mitochondrion sides (black arrows) suggestive of pre-fission events (Kleele *et al.*, 2021). Scale bar = 0.5 µm. D) Protein changes in the mitochondrial dynamics proteins FIS1 and MFN2 following MICE (n = 13) and SIE (n = 14). * denotes main effect at $p < 0.05$; # denotes time x group interaction at $p < 0.05$. For A), B), and

D) data are expressed as mean ± 95% confidence interval. For A) and B) differences were assessed with two-way ANOVA.

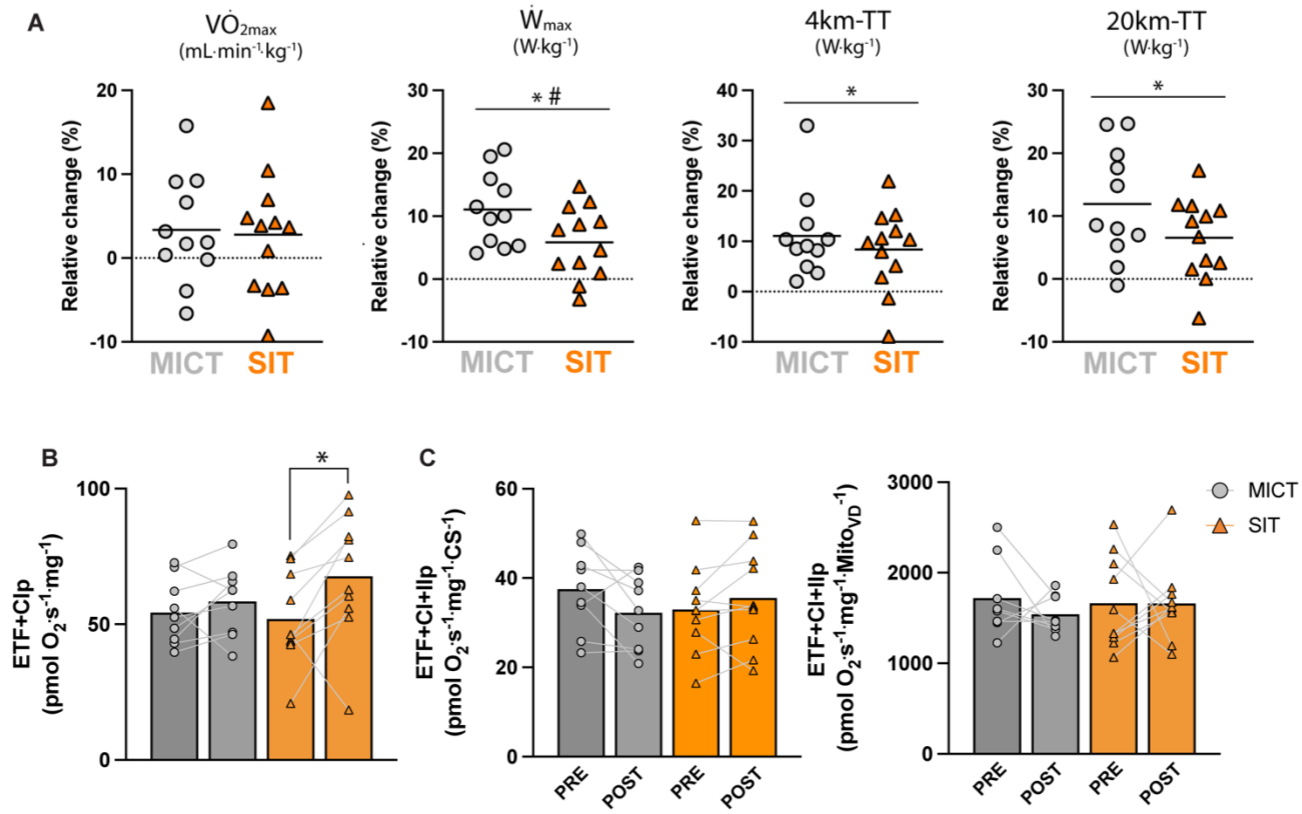


D

| | |
|---|---|
| Upregulated (MICT) | SLC25A3, NDUFS1, COX6C, ECH1, NDUFA11, SLC25A5, MGST3, NDUFA8, CS, NDUFB4, MT-CYB, ACO2, MRPS7, UQCRFS1, UQCRB, ACADVL, UQCRQ, NDUFB6, TUFM, ECI2, NDUFC2, NDUFB10, MRPL18, PRDX3, CHCHD3, NDUFB7, HSPA9, NDUFV1, ACSL1, MRPS36, HADHB, NDUFA7, COX7A1, FAM162A, NDUFA12, ETFB, GPT2, NDUFS3, UQCRC2, NDUFB9 |
| Upregulated (SIT) | LDHD, LONP1, GRPEL1, LYPLA1, AFG3L2, HINT3, AKR7A2, COMTD1, MRPL17, HSDL1, COASY, ACSS1, SFXN1, SIRT5, NAXD, ETHE1, HMGCL, MRPS27, SLC25A20, STOML2, RDH14, MRPL23, TRAP1 |
| Downregulated (MICT) | PDF, PMPCA, L2HGDH, FXN, C1QBP, MT-ND4, ACOT13, AK4, IDI1, NRDC, PTPMT1, PTRH2 |
| Downregulated (SIT) | SMDT1, NDUFS8, NDUFAF7, VDAC1, CYB5B, MRPL39, HINT2, SDHC, PAICS, COX6A2, TACO1 |

Supplemental Figure 5. A) Relative changes in maximal oxygen consumption ($VO_{2max}$), peak aerobic power ($W_{max}$), 4-kilometre time trial (4km-TT) performance, and 20-kilometre time trial (20km-TT) performance following moderate-intensity continuous training (MICT; n = 11) and sprint interval training (SIT; n = 12). B) Training-induced changes in respiratory function after addition of electron-transferring flavoprotein and complex I substrates (ETF+CIp). C) Training-induced mitochondrial-specific respiratory function (i.e., mass-specific respiration normalized to citrate synthase activity [MICT, n = 10; SIE, n = 11] or mitochondrial volume density [MICT, n = 9; SIE, n = 11]). D) List of the significant mitochondrial proteins differentially regulated following MICT and SIT in the proteomics data. * denotes main effect at $p < 0.05$; # denotes group x time interaction at $p < 0.05$. For A) and B) data are expressed as mean. For A) differences were assessed with two-way ANOVA. For B) differences were assessed with one-way ANOVA.

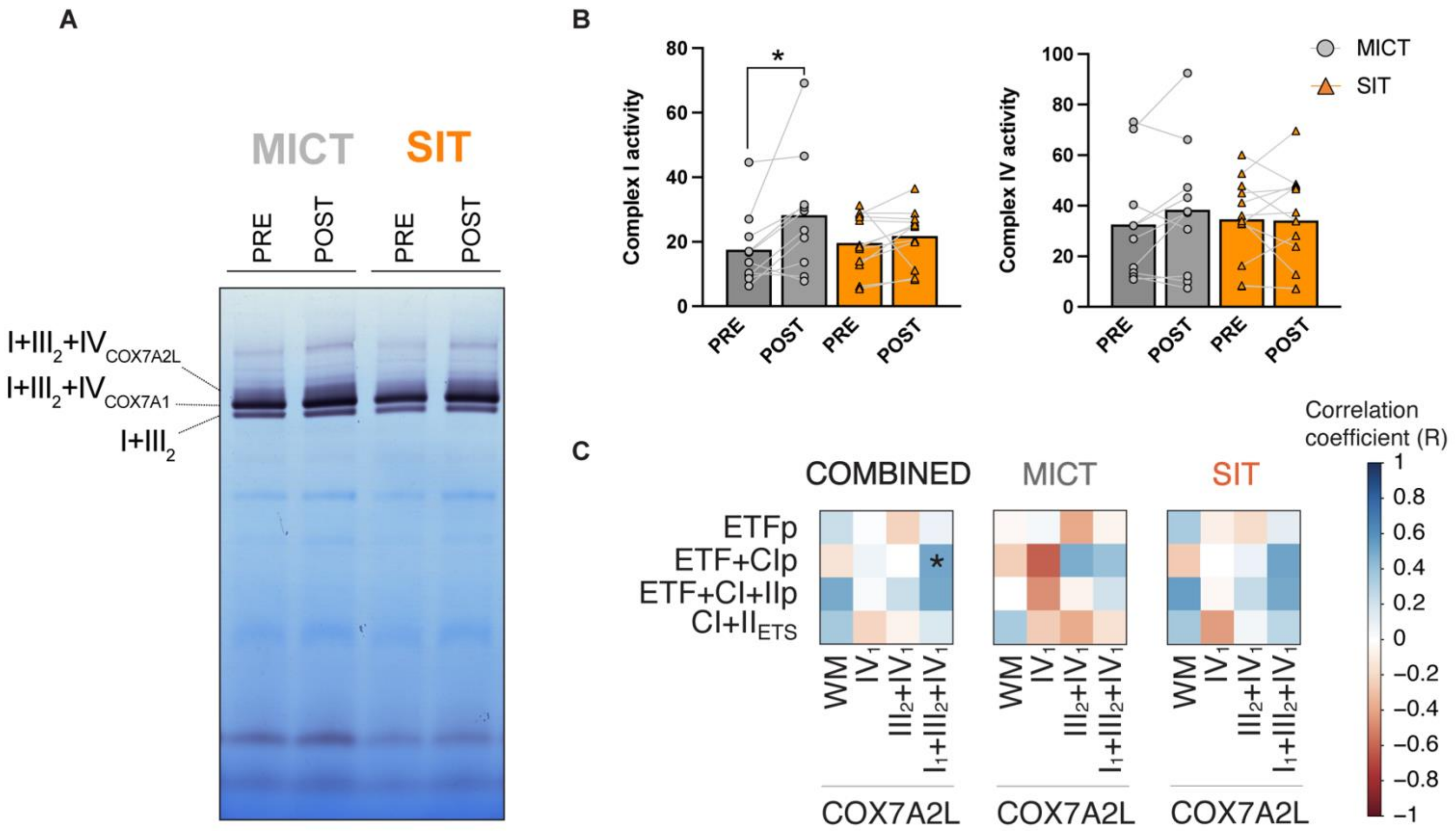


Supplemental Figure 6. A) Representative image of in-gel complex I enzymatic activity. B) Skeletal muscle complex I and IV activity in both moderate-intensity continuous training (MICT; n = 10) and sprint interval training (SIT; n = 12) groups. C) Pearson correlation of relative changes in mitochondrial respiratory function (ETFp = phosphorylation through electron-transferring flavoprotein [0.2 mM octanyolcarnitine, 2 mM malate, 3 mM MgCl2, 5 mM ADP]; ETF+CIp = phosphorylation through electron-transferring flavoprotein and complex I [0.2 mM octanyolcarnitine, 2 mM malate, 3 mM MgCl2, 5 mM ADP, 5 mM pyruvate]; ETF+CI+IIp = phosphorylation after addition of complex I- and complex II substrates [0.2 mM octanyolcarnitine, 2 mM malate, 3 mM MgCl2, 5 mM ADP, 5 mM pyruvate, 10 mM succinate]; $CI+II_{ETS}$ = maximal electron transport system capacity through complex I and II [obtained by 0.7-1.5 mM titrations of FCCP]) and relative changes in the protein content of COX7A2L measured in the whole muscle lysate (WM) or in different subcompartments ($IV_1$; $III_2$+IV1; $I_1+III_2+IV_1$). * = $p < 0.05$. For B) data are expressed as mean. For B) differences were assessed with pre-planned one-way ANOVA. For C) data was analyzed using Pearson correlation coefficient.
